# Alpha–beta oscillatory dynamics underlying auditory inhibitory control in children

**DOI:** 10.64898/2026.11.10.765597

**Authors:** Finnilä Elena, van Bijnen Sam, Kujala Jan, Parviainen Tiina

## Abstract

Response inhibition, the ability to withhold an unwanted response, is a fundamental component of cognitive control. While evoked responses in inhibitory tasks have been widely studied, induced oscillatory dynamics—particularly in the auditory modality and in children—remain largely unexplored. Here, we used magnetoencephalography (MEG) to examine induced alpha (8–12 Hz) and beta (13–20 Hz) oscillations during passive listening and an auditory Go/Nogo task in 67 children (6–14 years) and 16 adults (20–30 years). During passive listening, both groups showed an early alpha power increase in right fronto-temporo-parietal regions, followed by an alpha decrease in temporal regions. Under inhibitory demands, adults displayed a late alpha and beta power increase within a left fronto-temporo-occipital network, whereas children exhibited an early-onset, prolonged beta power decrease in somatomotor and temporo-parietal regions. Our findings indicate that children and adults engage spectrally, temporally, and spatially different neural dynamics during inhibitory control: in children, oscillatory modulation emerges earlier and is dominated by a beta decrease in sensorimotor regions, whereas adults recruit synchronized late alpha–beta dynamics in executive networks, possibly reflecting more efficient and evaluative inhibitory processing.

## 1. Introduction

Response inhibition is a core executive function that allows individuals to suppress automatic or inappropriate actions, and it is essential for flexible, goal-directed behavior (Aron, 2007; Bari & Robbins, 2013). This function encompasses both the filtering of sensory distractions and the halting of motor responses, and it develops substantially across childhood and adolescence, in parallel with the maturation of executive control systems (Luna et al., 2015; Tervo-Clemmens et al., 2023). Although response inhibition has been extensively studied through evoked brain responses, the role of oscillatory dynamics in inhibition—especially in children and in the auditory modality—remains underexplored. Endogenous oscillatory dynamics have been attributed a core role particularly in top-down–driven computations in brain information processing. Thus, this gap seriously limits our understanding of the neural basis underlying the emergence of cognitive control.

Building on animal models and clinical findings, response inhibition has been suggested to rely on a distributed network of cortical and subcortical structures. The cortico-subthalamic-pallidal circuit, incorporating both indirect and hyperdirect pathways, supports both sustained and rapid forms of motor inhibition (Nambu et al., 2002; DeLong & Wichmann, 2007). Prefrontal regions, including the inferior frontal gyrus (IFG), pre-supplementary motor area (pre-SMA), and anterior cingulate cortex (ACC), have been associated with coordination of top-down control, conflict monitoring, and error detection in a large review mainly drawing on functional magnetic resonance imaging (fMRI) findings (Diamond, 2013). In fMRI meta-analyses, emotional and motivational regulation have been linked with the amygdala (Hung et al., 2018) and the ventral striatum (Fakhraei et al., 2021), further shaping inhibitory control.

Developmentally, inhibitory control matures in parallel with the reorganization of large-scale brain networks (Kang et al., 2022). FMRI studies of blood-oxygen-level-dependent (BOLD) responses in inhibitory tasks show that younger children recruit more diffuse and bilateral activity across frontal and posterior cortices, whereas with age, activity becomes more spatially focused on regions including the left IFG and dorsolateral prefrontal cortex (DLPFC) (Durston et al., 2002; Tamm et al., 2002). The maturation of the anterior cingulate cortex (ACC) has been associated with improvements in error processing and the regulation of behavior across adolescence (Ordaz et al., 2013; Velanova et al., 2009). In parallel, increased network-level integration supports the development of goal-directed control (Marek et al., 2015). However, these studies rely on methods with limited temporal resolution, preventing a precise characterization of the unfolding neural dynamics.

In order to capture the temporal dynamics of different neural processes involved in response inhibition, it is necessary to utilize time-sensitive neuroimaging measures. Magnetoencephalography (MEG) allows non-invasive measurement of both stimulus-evoked responses and modulation of ongoing cortical rhythms with high temporal and spatial precision (Hämäläinen et al., 1993; Baillet, 2017). Most neural-level developmental studies of inhibition rely on evoked responses (Cheng et al., 2019; Jonkman et al., 2003; Ponton et al., 2000; van Bijnen et al., 2023), which are strictly phase-locked to stimulus onset and consequently dominated by stimulus-related characteristics rather than internally driven processes. In contrast, induced oscillatory modulation reflects neural signaling that is time-locked but not phase-locked to stimulus onset. Oscillatory modulations are typically assessed by examining changes in oscillatory power relative to baseline, and this modulation can reveal neural dynamics associated with attention, sensory gating, and motor preparation. While evoked responses primarily reflect exogenous, stimulus-driven processes, oscillatory activity is typically associated with endogenous, top-down dynamics, although it may also be modulated or entrained by external stimulation (Chen et al., 2012; Lakatos et al., 2019; Tallon-Baudry & Bertrand, 1999). This distinction between evoked responses and induced oscillatory activity is especially relevant in children, whose neural timing and response variability differ markedly from those of adults (Casey et al., 2005), and in whom the networks supporting cognitive, top-down control are still maturing. Consequently, oscillatory dynamics provide a valuable complementary perspective on the functional development of cognitive control.

Among the major cortical frequency bands, alpha (8–12 Hz) and beta (13–30 Hz) rhythms are particularly relevant for inhibitory control, reflecting complementary aspects of cognitive and motor regulation. Alpha rhythms are associated with top-down attentional control, acting as an inhibitory gating mechanism that suppresses task-irrelevant sensory input and supports selective attention, including auditory selective inhibition (Klimesch, 2012; Huang et al., 2014; Klimesch et al., 2007; Jensen & Mazaheri, 2010; Obleser & Weisz, 2012; Strauß et al., 2014). Beta-band modulation, in contrast, is associated with motor control and the regulation of action states during response inhibition (Engel & Fries, 2010; Swann et al., 2009; Huster et al., 2013). Importantly, beta oscillations are not functionally uniform across their frequency range, with lower beta frequencies (approximately 13–20 Hz) suggested to be involved in the maintenance and suppression of motor states (Engel & Fries, 2010; Kilavik et al., 2013). From a developmental perspective, the efficiency and specificity of alpha-band modulation improve with age, reflecting the maturation of cortical inhibitory systems (Güntekin et al., 2023), while sensorimotor beta dynamics show age-related changes in timing and strength, suggesting a gradual specialization of motor control networks (Gaetz et al., 2010; Trevarrow et al., 2019). Oscillatory mechanisms supporting inhibitory control have been most extensively characterized in the visual and somatosensory modalities, where modulation of alpha power is a well-established marker of cortical inhibition (López-Madrona et al., 2024; Strauß et al., 2014). By contrast, evidence from auditory contexts with respect to the joint involvement of alpha- and beta-band dynamics in children remains comparatively limited. Together, these findings highlight the potential value of examining alpha-and low-beta-band dynamics as developmental markers of emerging cognitive control.

In the present study, we investigated the role of cortical rhythms in auditory response inhibition to deepen the understanding of neural mechanisms underlying cognitive control in middle childhood. This developmental period is particularly important in the development of executive functions, including response inhibition. Alpha and beta oscillations are known as core cortical rhythms supporting inhibitory control, yet the developmental trajectory of auditory alpha- and beta-band dynamics remains poorly understood. Using source-level MEG estimates, we examined the spatiotemporal dynamics of alpha- and beta-band oscillations during passive sound presentation versus the same sounds under Go/Nogo task demands. Specifically, we asked whether sound-induced oscillatory patterns change under inhibitory requirements, and how these patterns differ in their spatial profile and temporal characteristics in children and adults.

## 2. Methods

### 2.1. Participants

A total of 94 participants, 78 children and 16 adults, took part in MEG recordings, structural MRI, and cognitive assessments (van Bijnen et al., 2023). All participants were Finnish-speaking and right-handed (as reported in a background questionnaire completed by the participants or, for children, by their parents). They were recruited from schools and the National Registry of Finland. None of the participants had neurological disorders or were on medication affecting the central nervous system, and their hearing was normal. Eleven children were excluded for the following reasons: one did not finish the experiment and one made too many errors in the MEG task (>50% errors in at least one block), five had extensive head movements or magnetic interference during MRI/MEG measurements, two did not agree to go into the MRI scanner, and two had structural abnormalities in their MRI. The final data included in this study consisted of 67 children (mean age: 10.2 years, SD: 1.4, range: 6‒14; 31 females, 36 males). All adult participants were retained in the analysis (mean age: 24.78, SD: 3.38, range: 20–30; 13 females, 3 males).

All participants underwent audiometric testing to confirm normal hearing. The study received approval from the Ethics Committee of the University of Jyväskylä, and informed consent was obtained from both children and their parents, following the principles of the Declaration of Helsinki. Compensation in the form of a movie ticket or gift card was provided to all participants for their involvement.

### 2.2. Stimuli, tasks, and procedure

The auditory stimuli consisted of 70 ms sine-wave tones presented at 65 dB SPL, with two frequencies: 1 kHz (standard tone, ST; 70% of trials) and 1.5 kHz (deviant tone, DT; 30%). Tones were generated using Audacity® (v2.3.3) and presented binaurally through an MEG-compatible high-fidelity sound system (Rotel RA-1570 amplifier) via silicone tubes and earpieces. Stimuli were delivered in a continuous stream with a variable interstimulus interval of 1.6–2.0 s. Each stream began with a ST, and consecutive DTs were avoided.

The participants completed three tasks using the same auditory stimuli: a passive listening task (PL), a Go/Nogo task (GN), and an oddball task (OB). In the PL task, participants were instructed to ignore the tones while watching a silent “Pingu” animation. In the GN task, they responded to STs; in the OB task, they responded to DTs. They responded with their right hand by pressing the outermost button of a four-button fiber-optic response pad (fORP, Current Designs Inc., Philadelphia, PA, USA) connected to the stimulus-presentation computer. The number of blocks and stimuli varied across tasks. There were two blocks of PL and four blocks each of GN and OB, with 150 trials per PL block (45 DTs, 105 STs) and 90 trials per GN and OB block (27 DTs, 63 STs). In the present report, we focus on the PL and GN DT conditions, neither of which required participants to make motor responses. The experiment was conducted in a child-friendly, submarine-themed game environment. Instructions were delivered by a virtual “captain” through video clips, and visual elements were designed by Studio Dennis Parren to enhance engagement. All stimuli were presented using PsychoPy (v3.2) running on Linux.

Before the tasks, participants completed four 1.5-minute resting-state recordings, alternating between eyes open and eyes closed conditions (two of each). Each task was preceded by 12 practice trials to ensure understanding. The session consisted of a PL block, followed by two alternating GN and OB blocks, a break, a second PL block, and two final GN and OB blocks presented in alternating order. During the break, participants were offered a snack and allowed to move around.

### 2.3. MEG and MRI recordings

Brain activity was recorded using a 306-channel magnetoencephalography (MEG) system with integrated electroencephalography (EEG) (Elekta Neuromag® TRIUX™, MEGIN Oy, Helsinki, Finland). The data were filtered between 0.1–330 Hz and sampled at 1000 Hz. Vertical and horizontal electro-oculograms (EOG) were used to detect eye movements and blinks for offline artifact correction. Electrodes were placed below and above the right eye and at the outer canthi of both eyes, with a ground electrode attached to the collarbone.

Five head position indicator (HPI) coils were attached to the EEG cap to track head position relative to the MEG sensor array throughout the session. The position of EEG electrodes and HPI coils was digitized relative to three anatomical landmarks (nasion, left and right preauricular points) using a Polhemus Isotrak digital tracker system (Polhemus, Colchester, VT, United States). Approximately 150 additional scalp points were digitized to enable co-registration with individual magnetic resonance imaging (MRI) data. Structural T1- and T2-weighted 3D spin-echo MRI images were acquired using a 1.5 T scanner (GoldSeal Signa HDxt, General Electric, Milwaukee, WI, USA) with a standard head coil and the following parameters: TR/TE = 540/10 ms, flip angle = 90°, matrix size = 256 × 256, slice thickness = 1.2 mm, sagittal orientation.

### 2.4. Preprocessing and source-level contrast analysis

The source-level analysis was conducted using event-related Dynamic Imaging of Coherent Sources (erDICS) beamforming (Laaksonen et al., 2012). Only MEG data were used for source modeling; EEG data were not included. To improve data quality, MaxFilter with movement-compensated temporal signal space separation (tSSS) was applied to reduce noise and correct head movements (Taulu & Simola, 2006), followed by independent component analysis (ICA) to remove artifacts such as eye blinks and heartbeats (Hyvärinen & Oja, 2000). The Go/Nogo deviant trials in which participants made a commission error (an incorrect button press to a deviant tone, operationalized as a press within 450 ms of deviant-tone onset) were identified and excluded from the MEG analysis prior to source estimation. On average, 8.3 (SD 7.0) trials were excluded in children and 5.2 (SD 3.6) in adults. The cross-spectral density (CSD) matrices for source modeling were computed using Morlet wavelets of width 7 in the range of 4–30 Hz. The CSD matrices were determined in the time window from –500 ms to 2,000 ms and downsampled by a factor of 5.

To correct for unequal trial counts between the passive and Go/Nogo deviant-tone conditions (90 vs. 108 presented trials), an equal number of trials per condition was randomly selected for each participant after excluding error trials (for the Go/Nogo task, only correct-rejection deviant trials), resulting in on average 89.7 (SD 2.4) trials per condition in children and 89.6 (SD 1.5) in adults. Next, oscillatory power in the alpha (8–12 Hz) and beta (13–20 Hz) bands at the source level was estimated using common beamformer weights across both experimental conditions and all examined time windows, with separate weights for the alpha and beta bands. The time windows for the analysis were set from 0 to 1,600 ms, each non-overlapping window having a duration of 200 ms. The forward model for source estimation was based on FreeSurfer MRI segmentation (Fischl, 2012) and was computed using MNE-Python (Gramfort et al., 2014). Prior to the statistical analysis, the individual-level estimates were transformed to a common coordinate system using a surface-based transformation in FreeSurfer (Fischl, 2012).

The modulation of oscillatory power in response to sounds was estimated for both PL and GN tasks relative to the baseline period (–400 ms to –200 ms before stimulus onset). Statistical significance was tested using four complementary contrasts: (1) one-sample t-tests assessed whether power modulations relative to baseline differed from zero, separately for PL and GN and for children and adults; all subsequent statistical contrasts were based on these baseline-corrected power values; (2) paired-samples t-tests examined task differences (PL vs. GN) in children and adults separately. In addition to these direct task-related effects within each age group, exploratory analyses were conducted to compare children and adults, using (3) independent-samples t-tests to compare groups (children vs. adults) within each task; and (4) independent-samples t-tests to assess group differences in task effects (PL vs. GN). Because of the unequal group sizes (67 children vs. 16 adults), contrasts (3) and (4) used Welch’s t-test, which does not assume equal variances between groups. All analyses were conducted in 200 ms time windows spanning 0–1,600 ms relative to stimulus onset. The significance level was set at p < .05, and to account for multiple comparisons, statistical results were corrected using the false discovery rate (FDR) method (Benjamini & Hochberg, 1995) within each time window and contrast. The resulting data were visualized using FreeSurfer 6.0.

#### 2.4.1. Region-of-interest time courses

To illustrate the power time courses underlying the whole-brain surface maps, we extracted ROI-averaged oscillatory power time courses (baseline-corrected; 200-ms windows, 100-ms steps) for three regions. ROIs were defined as complete Desikan–Killiany parcels, selected based on the strength of their baseline-related power modulation, quantified as the effect size (Cohen’s d) of the baseline-corrected power modulation, averaged in absolute value across both tasks (passive listening and Go/Nogo), both frequency bands (alpha, beta), and both groups (children and adults), and taking the maximum across time windows (see Supplementary Table S2). The three ROIs represent the two most prominently modulated, spatially distinct clusters in this ranking. All participants’ source estimates were interpolated to a common template source space, so that identical ROIs were used for children and adults. For each participant, task, and frequency band, power was averaged across the source points within each ROI. Within each ROI and frequency band, differences between tasks within each age group were tested with paired-samples t-tests, differences between age groups within each task with Welch’s t-tests, and the group × task interaction with Welch’s t-tests on the individual Go/Nogo *−* passive difference scores. For each of these comparisons, results were FDR-corrected (Benjamini–Hochberg, q < .05) across the time windows between 0 and 1,600 ms. Because ROIs were selected based on baseline-related modulation pooled across all tasks and groups, these comparisons are descriptive and are not independent of the ROI selection.

#### 2.4.2. Linear mixed-effects models

To complement the primary contrast-based analyses described above, we additionally modeled oscillatory power modulation across the entire cortical surface using linear mixed-effects (LME) models. For each cortical vertex, separately for each 200-ms time window (0–1,600 ms) and frequency band (alpha, beta), we fitted the model power ∼ task × age + (1 | subject), where task was a two-level categorical factor (passive listening vs. Go/Nogo, with passive listening as the reference level) and age (in years) was mean-centered across the sample. Subject was included as a random intercept to account for the within-subject, repeated-measures design. Four fixed-effect coefficients were extracted from each model: the intercept (mean baseline-corrected power during passive listening at the sample mean age), the main effect of task (the Go/Nogo-minus-passive difference at the sample mean age), the main effect of age (the linear age slope during passive listening, given the task × age interaction), and the task × age interaction (indexing whether the task effect changes with age, equivalently the linear age slope of the Go/Nogo-minus-passive difference score). As in the primary analyses, statistical significance was assessed using false discovery rate correction (Benjamini– Hochberg, q < .05), applied separately within each time window.

This model was fitted twice: first across the full sample (children and adults combined, N = 83; age mean-centered across the full sample), and second restricted to the children’s cohort only (N = 67, age range 6–14 years; age mean-centered within the children’s cohort). The children-only model serves as a robustness check, testing whether the task × age interaction observed in the full-sample model reflects a genuine, continuous age-related gradient rather than solely a between-group (child vs. adult) effect arising from the two age clusters in the full sample. Because the child and adult subsamples form two distinct, non-overlapping age clusters (6–14 and 20–30 years, respectively) rather than a continuously sampled age range, the full-sample age and task × age estimates are necessarily driven primarily by the between-group contrast and should not be interpreted as evidence for a smooth, continuous developmental trajectory across the unsampled 15–19-year interval. We therefore report these results as a complement to, rather than a replacement for, the categorical child–adult comparisons reported above. Full whole-brain results from both models are reported in the Supplementary Materials (Supplementary Figures S1–S4, full sample; S5–S8, children only).

### 2.5. Behavioral data analysis

In the Go/Nogo task, behavioral performance was evaluated based on three measures: reaction times (RTs) to standard tones, the intra-individual coefficient of variation (ICV), and the total number of errors. RTs and ICVs were calculated from correct responses only, with ICV defined as the standard deviation divided by the mean RT for each participant. Errors were defined as omissions (missed responses to standard tones) and commissions (false responses to deviant tones). Descriptive statistics for each age group are reported in Table 1. No inferential statistical analyses were conducted on behavioral data (van Bijnen et al., 2023).

**Table 1.**
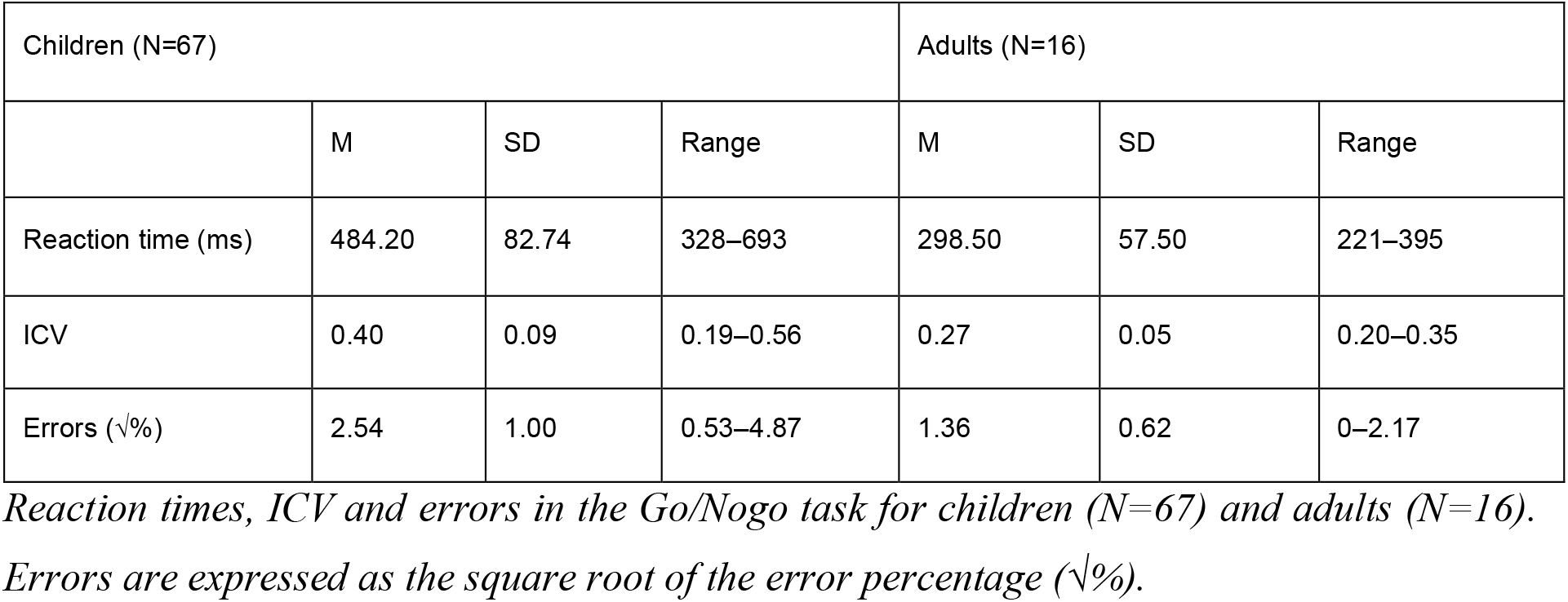
Behavioral performance in the auditory Go/Nogo task.

| Children (N=67) |  |  |  | Adults (N=16) |  |  |
|---|---|---|---|---|---|---|
|  | M | SD | Range | M | SD | Range |
| Reaction time (ms) | 484.20 | 82.74 | 328–693 | 298.50 | 57.50 | 221–395 |
| ICV | 0.40 | 0.09 | 0.19–0.56 | 0.27 | 0.05 | 0.20–0.35 |
| Errors (√%) | 2.54 | 1.00 | 0.53–4.87 | 1.36 | 0.62 | 0–2.17 |

## 3. Results

We examined the modulation of oscillatory power in the alpha and beta bands in response to simple sounds during passive listening and an inhibitory Go/Nogo task. Analyses were conducted separately for the two tasks and their contrasts in children and adults, both within and between age groups. These four analyses were designed to isolate task- and age-related components of oscillatory activity by systematically contrasting task effects within and across groups. This hierarchical approach helped to filter out non-specific modulations and highlight the core alpha- and beta-band dynamics most strongly associated with inhibitory demands. Full statistics for all significant clusters in the whole-brain contrasts (peak t-values, cluster sizes, and FDR-corrected p-values) are given in Supplementary Table S1.

### 3.1. Task-related modulation in oscillatory power relative to baseline in adults and children

#### 3.1.1. Passive listening task

In adults, during the passive listening task, a significant alpha power increase emerged in the right-lateralized fronto-temporo-parietal regions between 0 and 200 ms after stimulus presentation (peak t = 6.02), followed by significant alpha power decrease in the left auditory cortex between 400 and 600 ms (peak t = –6.18). No significant modulation of beta-band power was observed.

Children exhibited increased oscillatory power in both the alpha and beta bands in right-lateralized fronto-temporo-parietal regions between 0 and 200 ms after stimulus presentation (alpha: peak t = 4.57; beta: peak t = 5.76). This was followed by bilateral alpha power decrease from 200 to 1,000 ms (peak t = –7.96, 400–600 ms, left) and beta power decrease from 200 to 600 ms (peak t = –6.29, 200–400 ms, right). The decrease was most pronounced in temporal and inferior frontal regions. Figure 1 shows the distribution of regions with significant modulation of alpha and beta power relative to baseline during the passive listening task in both children and adults.

**Figure 1.**
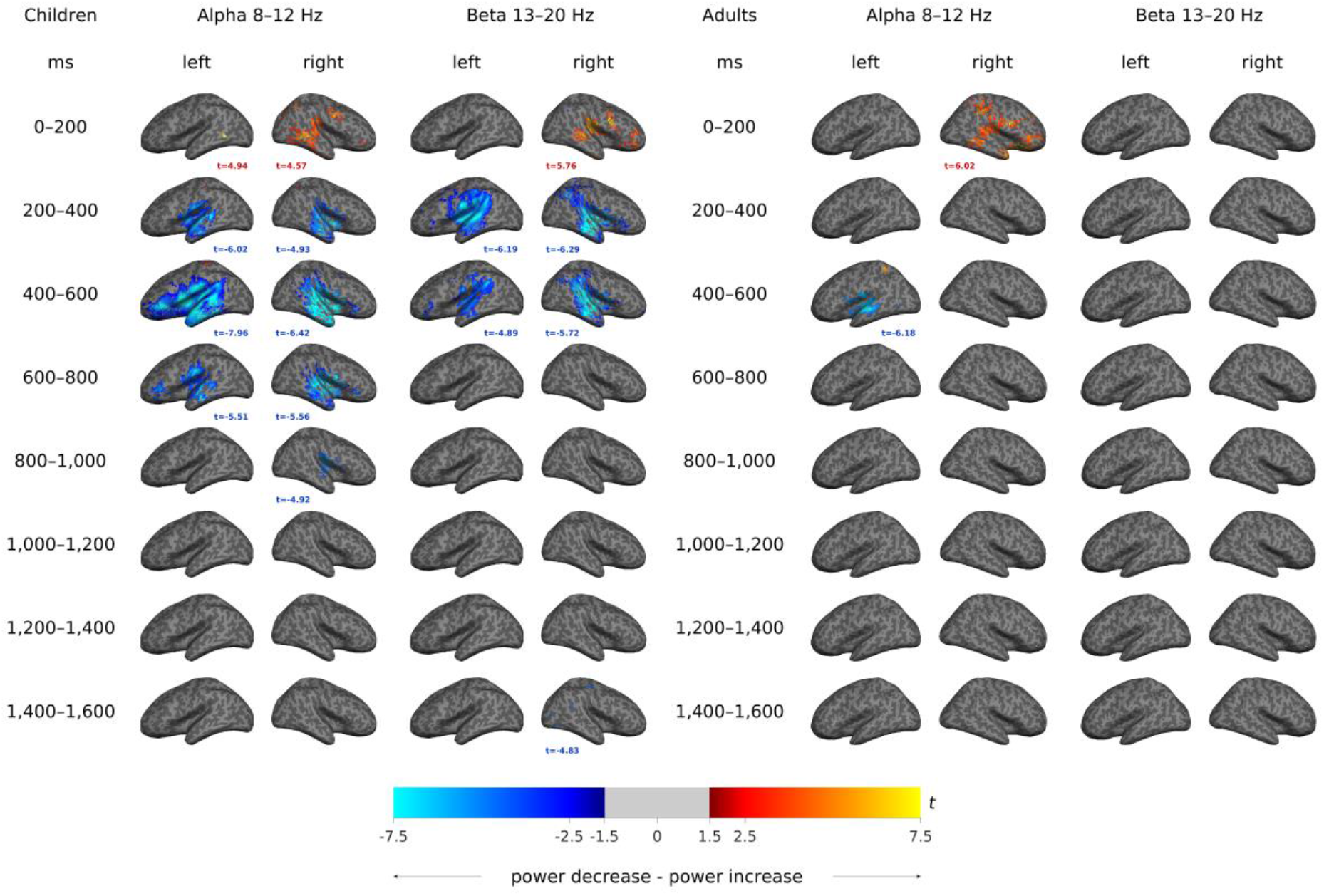
The passive listening task. Brain maps show sound-induced modulation of alpha (8– 12 Hz) and beta (13–20 Hz) power relative to baseline during the passive listening task in children and adults. Data are presented separately for the left and right hemispheres across consecutive time windows (0–200 ms, 200–400 ms, 400–600 ms, 600–800 ms, 800–1,000 ms, 1,000–1,200 ms, 1,200–1,400 ms, and 1,400–1,600 ms). Warm colors indicate power increases, and cool colors indicate power decrease. The peak t-value within each significant cluster (FDR-corrected, q < .05) is printed in the corresponding panel. Each brain surface shows a lateral view, with cortical regions including frontal, temporal, parietal, and occipital cortices.

#### 3.1.2. Go/Nogo task

In adults, there were no significant power modulations in the early time windows (0–400 ms).

A significant left-lateralized alpha power increase was observed between 400 and 1,000 ms after stimulus presentation (peak t = 5.73, left, 600–800 ms), and a beta power increase between 600 and 800 ms (peak t = 5.69, left). These modulations were distributed across temporal, frontal, and occipital regions.

In children, a significant bilateral alpha power increase was observed from 0 to 1,400 ms, initially widespread over somatomotor, posterior parietal, and prefrontal cortical regions (peak t = 6.69, left, 0–200 ms) and later confined to left prefrontal and parietal regions (peak t = 5.18, left, 1,000–1,200 ms). This power increase was overlapped by bilateral alpha power decrease between 200 and 1,000 ms in temporal regions (peak t = –6.07, left, 400–600 ms). Beta power showed strong, widespread bilateral decrease from 200 to 600 ms (peak t = –7.53, right, 200– 400 ms), continuing in the somatomotor areas up to 1,600 ms (peak t = –5.77, right, 1,200– 1,400 ms). Figure 2 shows the distribution of regions with significant modulation of alpha and beta power relative to baseline during the Go/Nogo task in both children and adults.

**Figure 2.**
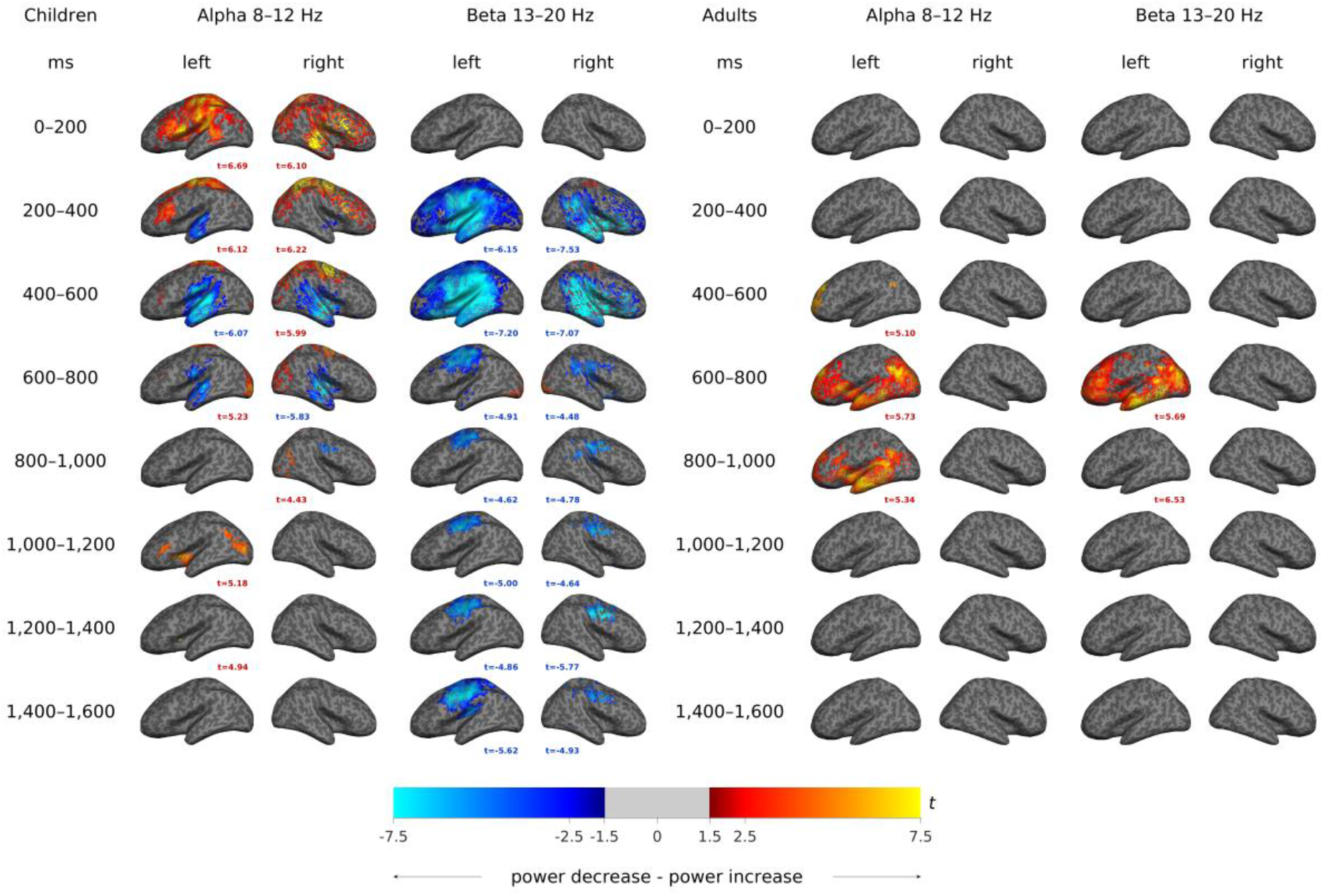
The Go/Nogo task. Brain maps show sound-induced modulation of alpha (8–12 Hz) and beta (13–20 Hz) power relative to baseline during the Go/Nogo task in children and adults. Data are presented separately for the left and right hemispheres across consecutive time windows (0–200 ms, 200–400 ms, 400–600 ms, 600–800 ms, 800–1,000 ms, 1,000–1,200 ms, 1,200–1,400 ms, and 1,400–1,600 ms). Warm colors indicate power increases, and cool colors indicate power decrease. The peak t-value within each significant cluster (FDR-corrected, q < .05) is printed in the corresponding panel. Each brain surface shows a lateral view, with cortical regions including frontal, temporal, parietal, and occipital cortices.

### 3.2. Differences in oscillatory power between task conditions (passive listening vs. Go/Nogo) in adults and children

In adults, differences between baseline-corrected oscillatory power during the passive listening and Go/Nogo tasks were observed in the alpha band from 400 to 1,000 ms, particularly over the left frontal, temporal, and occipital cortices, indicating consistently higher alpha power in the Go/Nogo task than in the passive task throughout this interval (peak t = 8.43, left, 600–800 ms). In the beta band, left-lateralized task-related differences were found between 400 and 800 ms, most prominently between 600 and 800 ms over frontal, temporal, and occipital regions, indicating a stronger beta power increase in the Go/Nogo task (peak t = 8.11, 600–800 ms).

In children, task differences in alpha power were observed bilaterally from 200 to 800 ms involving fronto-temporal regions, indicating a greater alpha power decrease in the passive task (peak t = 5.84, left, 200–400 ms). In addition, minor right-hemisphere clusters of the opposite sign (lower alpha power in the Go/Nogo task) were observed from 1,000 to 1,400 ms (peak t = –4.93). In the beta band, task differences were observed between 200 and 1,000 ms and from 1,400 to 1,600 ms in the left parietal, temporal, and somatomotor regions, indicating a stronger beta power decrease in the Go/Nogo task (peak t = –4.93, left, 600–800 ms), with small right-hemisphere clusters from 200 to 400 ms and from 1,200 to 1,400 ms (peak t = –5.04, 1,200– 1,400 ms). Figure 3 shows the distribution of regions with significant task differences in alpha- and beta-band power modulation relative to baseline in both children and adults.

**Figure 3.**
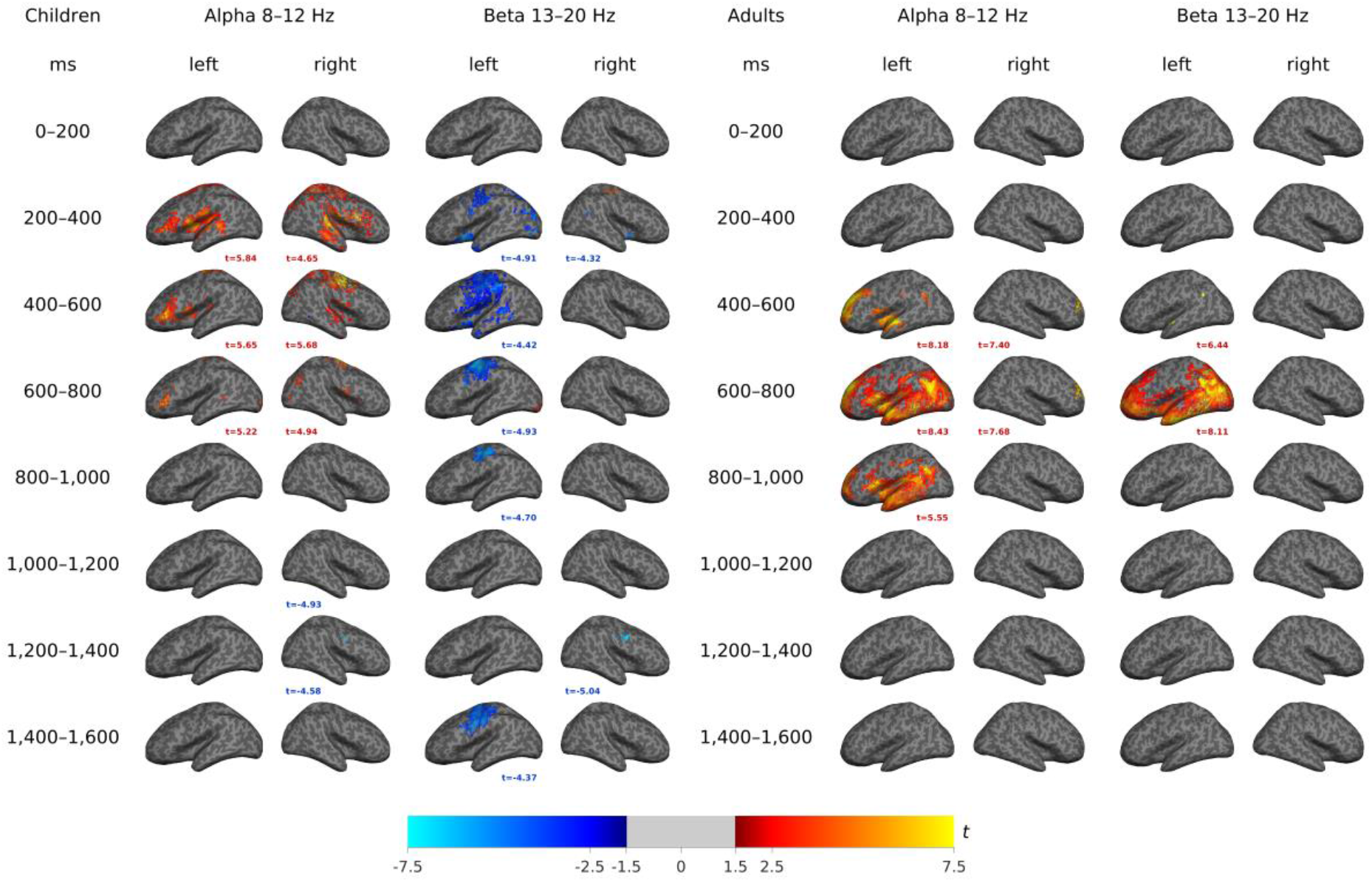
Passive listening vs. Go/Nogo. Brain maps show differences in sound-induced modulation of alpha (8–12 Hz) and beta (13–20 Hz) power between the passive listening and Go/Nogo tasks in children and adults. Data are presented separately for the left and right hemispheres across consecutive time windows (0–200 ms, 200–400 ms, 400–600 ms, 600–800 ms, 800–1,000 ms, 1,000–1,200 ms, 1,200–1,400 ms, and 1,400–1,600 ms). Warm colors indicate higher baseline-corrected power during the Go/Nogo task than during passive listening (Go/Nogo – passive > 0), and cool colors the reverse. The peak t-value within each significant cluster (FDR-corrected, q < .05) is printed in the corresponding panel. Each brain surface shows a lateral view, with cortical regions including frontal, temporal, parietal, and occipital cortices.

### 3.3. Differences in oscillatory power between age groups in the passive listening and Go/Nogo tasks

We then analyzed differences between adults and children separately for the two tasks (examining modulation of oscillatory power relative to baseline during the passive listening and Go/Nogo tasks contrasted between age groups), as well as for the task-contrast (the difference in baseline-corrected oscillatory power between the passive listening and Go/Nogo tasks contrasted between age groups).

#### 3.3.1. Passive listening task

No significant differences were found between children and adults during the passive listening task (Figure 4).

**Figure 4.**
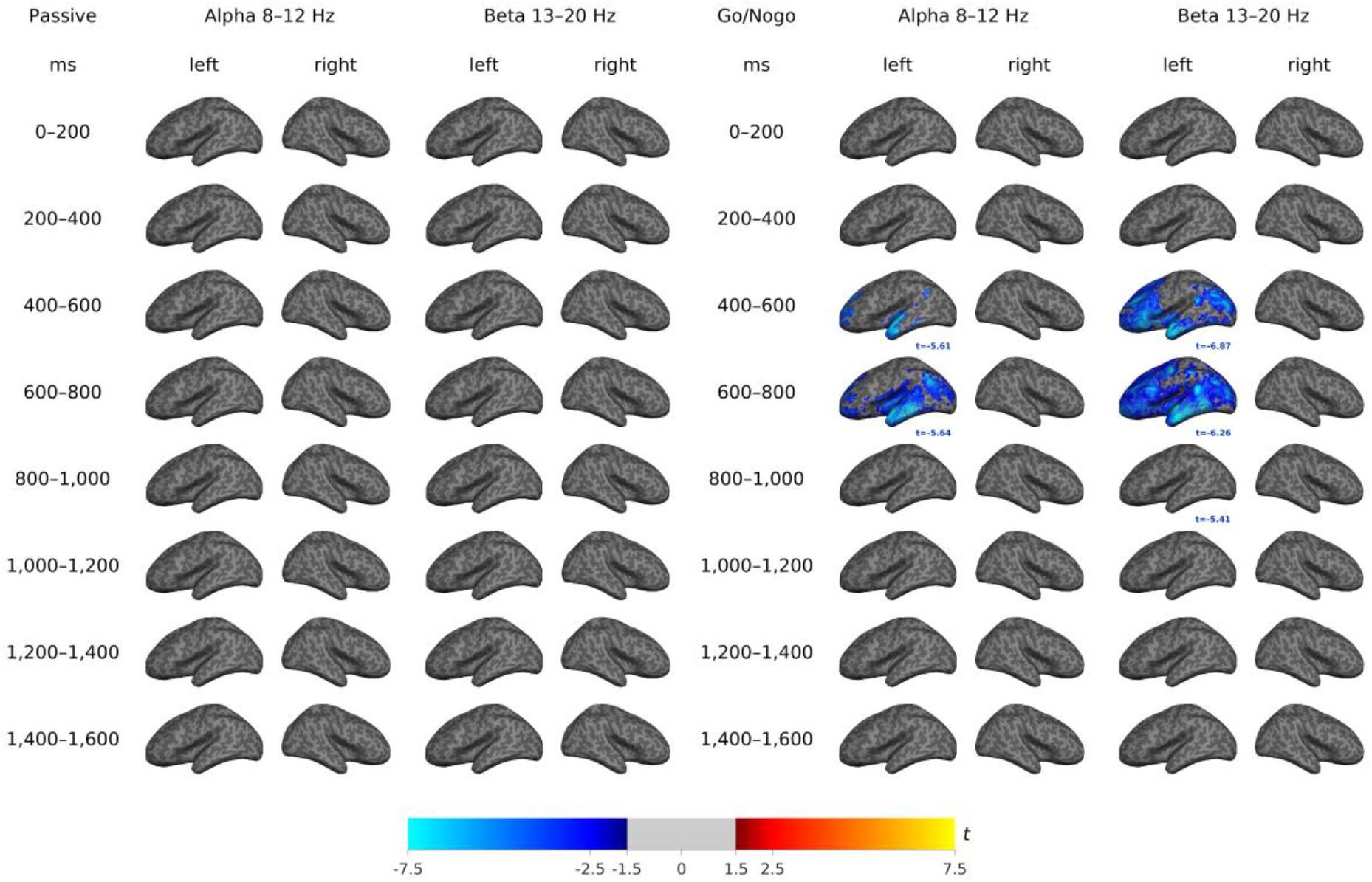
Age-group differences during the passive listening and Go/Nogo tasks. Brain maps show differences in sound-induced modulation of alpha (8–12 Hz) and beta (13–20 Hz) power between children and adults during the passive listening task and Go/Nogo task; no significant differences were found during passive listening. Data are presented separately for the left and right hemispheres across consecutive time windows (0–200 ms, 200–400 ms, 400–600 ms, 600–800 ms, 800–1,000 ms, 1,000–1,200 ms, 1,200–1,400 ms, and 1,400–1,600 ms). Warm and cool colors indicate the direction of group differences, calculated by subtracting the oscillatory modulation of adults from that of children. The peak t-value within each significant cluster (FDR-corrected, q < .05) is printed in the corresponding panel. Each brain surface shows a lateral view, with cortical regions including frontal, temporal, parietal, and occipital cortices.

#### 3.3.2. Go/Nogo task

Children showed significantly different oscillatory power modulation than adults during the Go/Nogo task, confined to the left hemisphere between 400 and 1,000 ms after stimulus presentation, involving frontal, temporal, and parietal regions. In the alpha band, significant group differences were observed from 400 to 800 ms (peak t = –5.64, 600–800 ms); in the beta band, from 400 to 1,000 ms (peak t = –6.87, 400–600 ms). All significant clusters were left-lateralized and uniformly reflected a smaller power increase (or a larger decrease) in children relative to adults. Figure 4 shows the distribution of regions with significant differences in alpha and beta power between children and adults during the Go/Nogo task.

#### 3.3.3. Passive listening vs. Go/Nogo task

The between-group difference in the task contrast (passive listening vs. Go/Nogo) was likewise confined to the left hemisphere, between 400 and 800 ms, involving frontal, temporal, and parietal regions. Significant differences were observed in the alpha band (peak t = –5.62, 600– 800 ms), with a relatively small and spatially restricted cluster, and in the beta band (peak t = –6.03, 600–800 ms), which showed a larger and more spatially extensive cluster over the same regions, both reflecting a relatively smaller alpha/beta power increase (or a larger decrease) in children’s task-related modulation compared with adults’. Figure 5 shows the distribution of regions with significant differences in alpha and beta power in the between-group contrasts of task effects.

**Figure 5.**
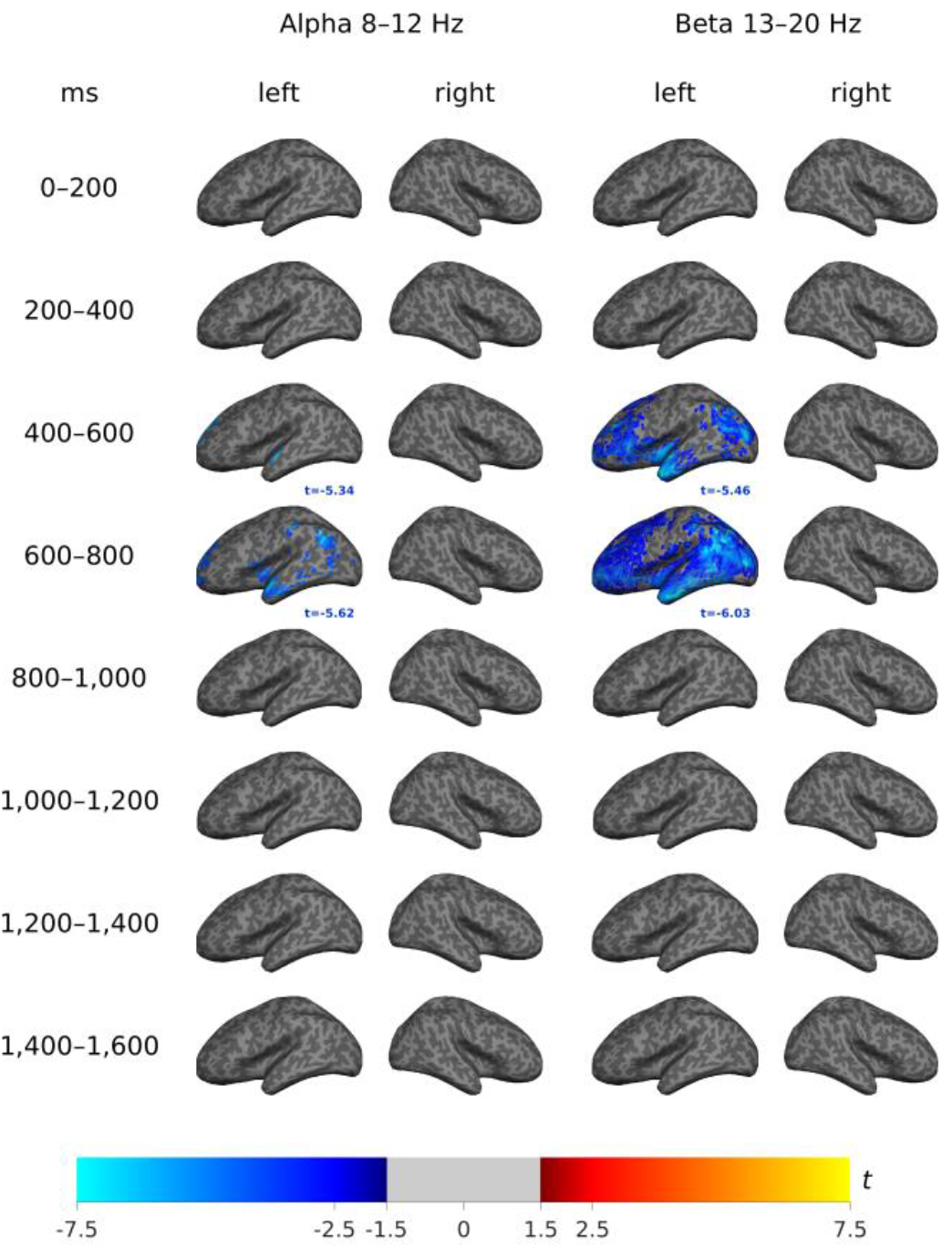
Differences between children and adults in the passive listening vs. Go/Nogo task contrast. Brain maps show differences in sound-induced modulation of alpha (8–12 Hz) and beta (13–20 Hz) power between children and adults for the task contrast (passive listening vs. Go/Nogo). Data are presented separately for the left and right hemispheres across consecutive time windows (0–200 ms, 200–400 ms, 400–600 ms, 600–800 ms, 800–1,000 ms, 1,000–1,200 ms, 1,200–1,400 ms, and 1,400–1,600 ms). Warm and cool colors indicate between-group task differences, calculated by subtracting the difference between tasks in oscillatory modulation of adults from that of children. The peak t-value within each significant cluster (FDR-corrected, q < .05) is printed in the corresponding panel.

#### 3.3.4. Region-of-interest time courses

The most prominent power modulation was observed bilaterally in the temporal cortex (superior, transverse, and middle temporal gyri). The insula, the supramarginal gyrus, and the parahippocampal cortex, which were also highly ranked, lie adjacent or close to this temporal cluster and likely reflect spatial spread (beamformer leakage) from the same underlying source. The next spatially distinct cluster of modulation was observed in the left inferior frontal cortex, comprising pars triangularis, pars orbitalis, and pars opercularis, regions with an established role in inhibitory control. The complete list of cortical parcels, ranked by overall power modulation relative to baseline, is shown in Supplementary Table S2. To further illustrate these effects, we selected three ROIs representing these two spatially distinct clusters for time-course analysis: the left and right lateral temporal cortex and the left inferior frontal gyrus (IFG; Figure 6). Within each ROI, we compared the two tasks within each age group, the two age groups within each task, and the task effect between groups (t-values: Go/Nogo – passive for task effects, children – adults for group effects); these comparisons are descriptive.

**Figure 6.**
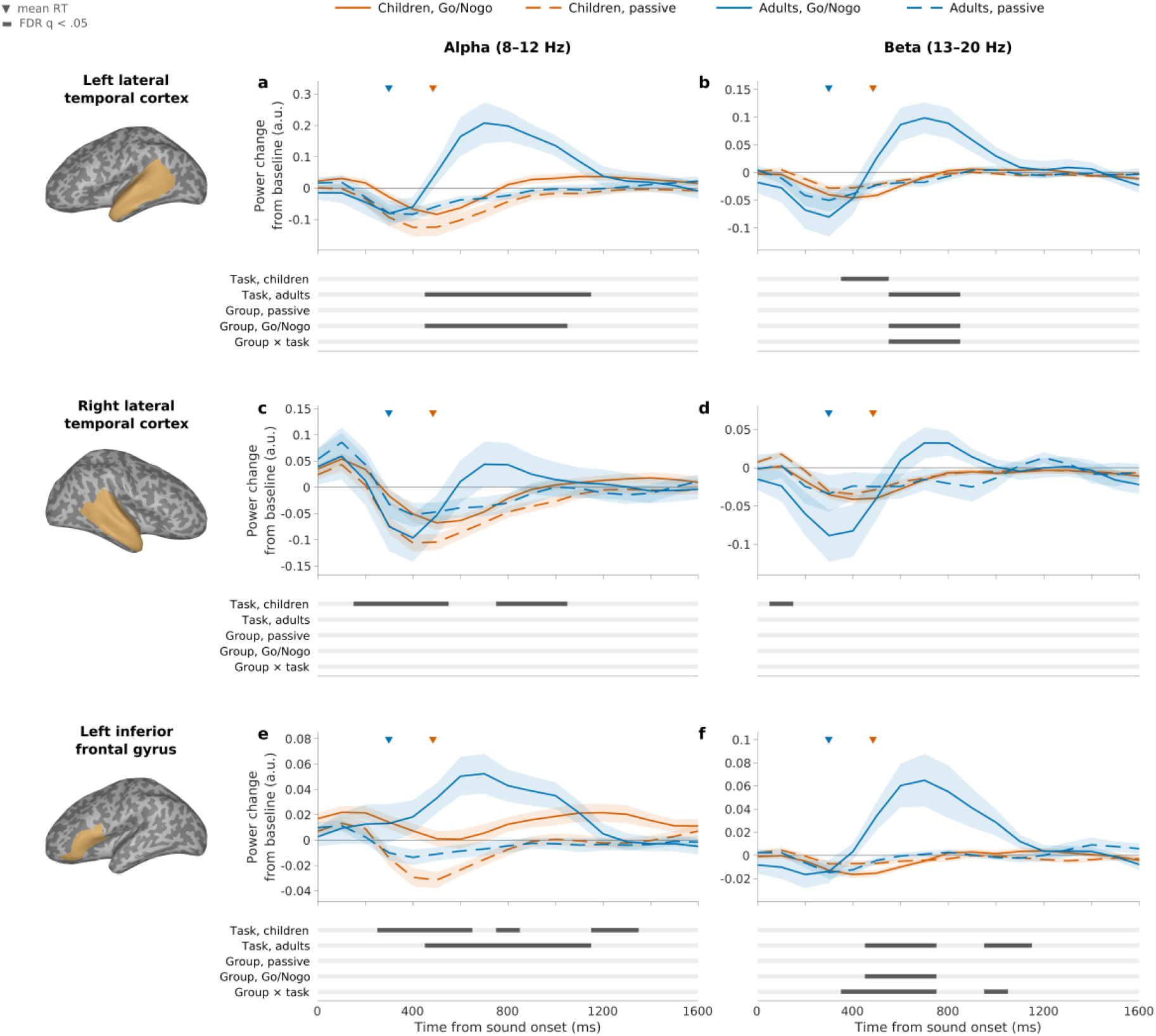
ROI-averaged alpha and beta power time courses. Baseline-corrected alpha (8–12 Hz; left column) and beta (13–20 Hz; right column) power in the left lateral temporal cortex (a, b), the right lateral temporal cortex (c, d), and the left inferior frontal gyrus (IFG; e, f) in children (orange; n = 67) and adults (blue; n = 16) during the Go/Nogo task (solid lines) and passive listening (dashed lines). ROIs were defined as complete Desikan–Killiany parcels (selection based on Supplementary Table S2): lateral temporal cortex = superior temporal, transverse temporal, and middle temporal gyri and the banks of the superior temporal sulcus; IFG = pars opercularis, pars triangularis, and pars orbitalis. They are shown on the inflated cortical surface of the template participant. Power was averaged across all source points within each ROI and expressed as the change from the baseline window (−400 to −200 ms). Values are plotted at the centers of 200-ms windows (100-ms steps); shading indicates ±SEM. Triangles mark the mean Go-trial reaction time of each group. Bars below each panel indicate time windows with significant differences (Benjamini–Hochberg FDR-corrected across time windows, q < .05) between tasks within each group (Task; paired t-tests), between groups within each task (Group; Welch’s t-tests), and in the task effect between groups (Group × task; Welch’s t-tests on the Go/Nogo − passive differences). These comparisons are descriptive. Note the different y-axis scales across panels.

In the left lateral temporal cortex, adults showed higher alpha power during the Go/Nogo task than during passive listening (400–1,200 ms; peak t = 4.54), and during the Go/Nogo task, alpha power was higher in adults than in children (400–1,100 ms; peak t = –3.65). A similar but shorter pattern was observed in beta power (500–900 ms): the task effect in adults (peak t = 3.98) was accompanied by a group difference during the Go/Nogo task (peak t = –3.78) and a group × task interaction (peak t = –3.91). In children, beta power in this region was lower during the Go/Nogo task than during passive listening between 300 and 600 ms (peak t = – 3.51). In the right lateral temporal cortex, the Go/Nogo-related increase in adults was weaker and not significant, whereas children showed higher alpha power during the Go/Nogo task than during passive listening (100–600 ms, peak t = 4.25; 700–1,100 ms, peak t = 2.90), reflecting a smaller alpha decrease during the Go/Nogo task.

In the left IFG, alpha power was higher during the Go/Nogo task than during passive listening in both age groups: in adults from 400 to 1,200 ms (peak t = 3.62), and in children in three shorter intervals between 200 and 1,400 ms (peak t = 3.26). In beta power, a task effect was observed only in adults (400–800 ms, peak t = 3.23; 900–1,200 ms, peak t = 3.86), resulting in a group difference during the Go/Nogo task (400–800 ms; peak t = –3.98) and a group × task interaction (300–800 ms, peak t = –3.47; 900–1,100 ms, peak t = –2.67). No group differences were observed during passive listening in any of these ROIs.

Together, these time courses illustrate that, in adults, Go/Nogo-related modulation was characterized by a relatively synchronized increase in alpha and beta power in the left temporal cortex and left IFG at approximately 500–1,000 ms, which was descriptively stronger in the left than in the right temporal cortex. This increase was absent in children; instead, their task effects consisted of a smaller alpha decrease during the Go/Nogo task than during passive listening in the right temporal cortex and the left IFG, and a brief beta decrease in the left temporal cortex.

### 3.4. Linear mixed-effects analysis

To complement the categorical task- and age-group contrasts reported above, we additionally examined oscillatory power modulation using a vertex-wise linear mixed-effects model incorporating task, continuous age, and their interaction, fitted twice: across the full sample (N = 83) and restricted to the children’s cohort alone (N = 67, age range 6–14 years). Full whole-brain results are reported in the Supplementary Materials (Supplementary Figures S1–S4, full sample; S5–S8, children only).

In the full sample, the model intercept (mean baseline-corrected power during passive listening at the sample mean age) closely mirrored the categorical passive-listening results reported in Section 3.1.1: an early (0–200 ms) alpha and beta power increase in the right hemisphere (peak t = 5.60 and t = 4.91), followed by a widespread bilateral alpha and beta power decrease peaking at 400–600 ms for alpha (peak t = –7.72) and at 200–400 ms for beta (peak t = –6.38), and resolving after 1,000 ms (Supplementary Figure S1). The main effect of task (Go/Nogo minus passive) likewise converged with the categorical task-contrast findings (Section 3.2): a robust, bilateral alpha power increase emerged predominantly between 200 and 1,000 ms (peak t = 6.53), while beta power showed an early decrease (0–400 ms, peak t = – 4.68), a marked increase between 400 and 800 ms (peak t = 5.24), and a further decrease from 1,000 to 1,600 ms (Supplementary Figure S2).

In contrast, the main effect of age (the linear age slope during passive listening) was essentially absent in the full sample, with only a single small cluster surviving FDR correction (right hemisphere beta, 800–1,000 ms, peak t = 4.23; Supplementary Figure S3). This null result is consistent with the categorical finding that children and adults did not differ during passive listening (Section 3.3.1).

The task × age interaction, by contrast, was the most extensive effect in the full-sample model: a widespread bilateral beta power increase spanning nearly the entire lateral cortical surface was observed between 400 and 1,000 ms (peak t = 7.71, 600–800 ms), accompanied by a more spatially restricted, predominantly left-lateralized alpha power increase over the same interval (peak t = 6.22), together with smaller beta power decreases in sensorimotor regions before 400 ms and after 1,200 ms (Supplementary Figure S4). This interaction likely reflects the divergent task-related beta modulation between children, who showed a power decrease, and adults, who showed a power increase, during this interval (Sections 3.1.2 and 3.2).

Critically, when the same task × age interaction was estimated within the children’s cohort alone, where age is sampled continuously rather than forming two discrete clusters, only two small clusters survived FDR correction (both right hemisphere beta: 200–400 ms, peak t = – 4.66; 600–800 ms, peak t = 4.65; Supplementary Figure S8). The later, positive cluster coincided with the time window in which adults show their characteristic beta power increase (Section 3.1.2), which is tentatively consistent with an emerging, age-related shift toward the adult-typical pattern. The corresponding main effect of age within children was similarly sparse (one cluster, right hemisphere beta, 400–600 ms, peak t = 4.45; Supplementary Figure S7). As anticipated in Section 2.4.2, this indicates that the extensive full-sample task × age interaction is driven primarily by the categorical difference between the child and adult subsamples rather than by a continuous developmental gradient within the 6–14-year age range sampled here, consistent with the categorical finding that the largest group differences emerged specifically during the Go/Nogo task (Section 3.3.2). The intercept and task-effect models within the children’s cohort (Supplementary Figures S5–S6) converged closely with both the full-sample model and the categorical within-group findings reported in Sections 3.1.1 and 3.2, providing additional, model-based support for the categorical contrasts that form the primary analyses of this study.

## 4. Discussion

This study examined the role of alpha and beta oscillations in response inhibition in mid-childhood. Using source-level analysis of MEG data, we found that although auditory stimulation during passive listening induced largely similar spatial and temporal patterns of oscillatory power in children and adults, distinct spectral, temporal, and spatial patterns emerged when the task required inhibitory control.

### 4.1. Auditory processing during passive listening

During passive listening, children exhibited an early alpha power increase in right-lateralized fronto-temporo-parietal regions, followed by bilateral alpha power decrease in the temporal and inferior frontal regions. There were no significant differences compared to adults, who likewise showed an early alpha power increase in right-lateralized fronto-temporo-parietal regions and subsequent alpha power decrease in the left temporal cortex (Figure 1). These findings align with previous reports of auditory processing and likely reflect established mechanisms underlying cortical responses to simple sounds (Weisz et al., 2011). Consistent with this, both MEG and intracranial recordings have shown stimulus-induced alpha power decrease in the auditory cortex during passive listening in adults (López-Madrona et al., 2024) and in children (Fujioka & Ross, 2008). Studies using evoked potentials have revealed developmental differences in auditory-evoked responses, such as an emphasis on the late-evoked N250 component in children (Parviainen et al., 2019), suggesting that immature phase-locked auditory processing is associated with prolonged processing periods (Cheng et al., 2019; Ponton et al., 2000; Ruhnau et al., 2011). Interestingly, our findings indicate that induced oscillatory responses in children do not differ from those in adults during the passive listening task, suggesting that developmental differences in oscillatory modulation may emerge primarily under higher cognitive demands.

### 4.2. Inhibitory control

The Go/Nogo task was associated with distinct oscillatory dynamics underlying inhibitory control in children and adults. In adults, alpha and beta power increased within a left-lateralized fronto-temporo-occipital network, most prominently from approximately 600 ms after stimulus onset (and 300 ms after the anticipated (and inhibited) response based on go-trial response times). Relative to baseline, children showed a bilateral alpha power increase overlapped by prolonged decrease in alpha power and even stronger and more widespread decrease in beta power across cortical regions, extending throughout the period of sound processing and the anticipated (inhibited) response, as estimated based on go-trial response times. Direct comparisons between the tasks and age groups further clarified the use of cortical resources for response inhibition (Figures 3–5). In adults, differences between the tasks showed relatively increased alpha–beta power for Nogo trials compared to passive listening, with alpha decrease present during passive listening but absent in the Go/Nogo task. In children, task-related differences were observed in both frequency bands: a more temporally extended decrease of beta power during Nogo trials compared to passive listening, accompanied by a spatially more extensive but temporally more restricted alpha-band difference, reflecting a greater alpha power decrease during passive listening than during the Go/Nogo task. Group contrasts during Nogo trials (Figure 4) and between-group contrasts of task effects (Figure 5) were confined to the left frontal, temporal, and parietal cortices between 400 and 1,000 ms. In both, children showed a smaller power increase (or a larger decrease) than adults. Together with the within-group results (Figures 2 and 3), this indicates that adults display late alpha and beta power increases in left-lateralized frontal, temporal, and occipital cortices. Children, by contrast, exhibit an early emerging and prolonged beta power decrease during the Go/Nogo task, which is bilateral but most extensive in the left somatomotor and temporo-parietal regions. Overall, these findings indicate that inhibitory demands engage distinct neural patterns in children and adults, differing in *spatial distribution, timing, duration, and the direction* of oscillatory power modulation (decrease vs. increase).

### 4.3. From diffuse decrease to coordinated oscillatory control

Interestingly, inhibitory demands *increased* oscillatory power in adults, whereas in children they elicited a pronounced decrease. This basic divergence likely reflects differences in underlying neural processes as well as in the efficiency of audio-motor integration and response times: in children, this decrease may reflect slower or more elaborated recruitment of the motor system, whereas in adults, the enhanced rhythmic synchronization may reflect more efficient top-down control or evaluative processes following successful inhibition (cf. next paragraphs). At the spectral level, children’s responses were dominated by a decrease in the beta band.

In contrast, adults exhibited coordinated increases in both alpha and beta power. This coordinated alpha–beta pattern in adults may reflect more integrated executive functioning, allowing for flexible adjustment of both sensory gating and motor output (Horn et al., 2003; Swann et al., 2009). In children, the reliance on beta power decrease may reflect a more reactive, motor-focused strategy during the time window when the response is expected (based on children’s go-trial reaction times). This interpretation aligns with previous findings associating alpha oscillations with top-down suppression of task-irrelevant information (Klimesch, 2012; Huang et al., 2014) and beta rhythms with motor control, particularly the maintenance or termination of ongoing task-related processes (Cavanagh & Frank, 2014; Engel & Fries, 2010; Swann et al., 2009).

Temporally, the groups diverged markedly. In children, the beta decrease emerged early, beginning around 200 ms post-stimulus, prior to the average reaction time (484.20 ms; Figures 2 and 6), and persisted in somatomotor regions well beyond the behavioral response, continuing into the later stages of the processing window (Figure 2). This early and sustained modulation likely reflects a preparatory or diffuse motor readiness state, suggesting that children engage sensorimotor systems proactively, perhaps due to immature inhibitory processing. Adults, by contrast, showed temporally focused increases in oscillatory power that emerged later, from approximately 400–600 ms, and peaked at 600–800 ms (Figures 2, 3, and 6). Because adults typically respond to standard tones within 200–400 ms, this late activity may reflect a post-inhibitory rebound or evaluative process following successful inhibition. Alternatively, it may correspond directly to the inhibitory process itself, which typically has a longer latency than standard motor responses because it requires stimulus evaluation and the cancellation of a prepared action. Studies focusing on evoked responses show that in adults, the core decision point of the inhibition process appears earlier (the 200–300 ms time window) than the induced oscillatory changes observed here, thus our oscillatory modulation is likely related to the stabilization, maintenance, or evaluation of inhibitory control across a more extended temporal window (van Bijnen et al., 2023). Together, these findings suggest that in the mature brain, response inhibition involves phase-locked, stimulus-evoked activity that is accompanied by later oscillatory dynamics that may reflect evaluative processes. Although the within-children LME analyses indicated that most of this temporal divergence reflects the categorical child– adult gap rather than a continuous gradient, the later of the two small clusters that survived within the children’s cohort fell within this same later time window and pointed in the adult-typical direction, albeit in the right rather than the left hemisphere. This hints that the shift toward this late oscillatory pattern may already be beginning to emerge, even if subtly, within middle childhood.

The spatial distribution of oscillatory responses further differentiates children and adults. Children showed the strongest effects in the somatomotor and temporo-parietal regions, suggesting a reliance on sensorimotor circuits. Adults, on the other hand, recruited a left-lateralized fronto-temporo-occipital network. The left lateralization of activation aligns with evoked activity reported from the same dataset, where the amplitude of the N250m response in the left hemisphere uniquely predicted behavioral variability in the inhibition task (ICV, errors, SSRT in auditory Go/Nogo task) (van Bijnen et al., 2023). These findings imply that both the evoked and oscillatory activity in the left hemisphere play a functionally central role in inhibitory control. The engagement of frontal areas in adults aligns with the findings on the involvement of higher-order executive regions in mature inhibitory control (Rubia et al., 2007; Huster et al., 2013). In contrast, the absence of frontal involvement in children may reflect immature integration of executive control networks, consistent with a developmental account proposing a gradual shift from effortful, executive-driven inhibition toward more automatized control processes (Diamond, 2013). However, it may also be that a similar fronto-temporo-occipital network would also emerge in children only with longer inter-stimulus intervals.

Taken together, these spectral, temporal, and spatial differences suggest a developmental shift from broad, prolonged decrease in motor areas to temporally precise and spatially coordinated control involving a fronto-temporo-occipital executive network. In this framework, children’s neural dynamics may reflect a more effortful, diffuse inhibitory strategy, whereas adults engage more automatic, targeted mechanisms; possibly relying on learned task structures and internalized control schemas. This interpretation is consistent with behavioral findings, including longer and more variable reaction times in children, and supports the notion that efficient response inhibition may depend on the maturation of synchronized alpha–beta dynamics in distributed control networks.

### 4.4. Implications for cognitive control development

Our findings contribute to the growing body of literature highlighting the role of oscillatory dynamics in inhibitory control across development. These developmental patterns have important implications for understanding atypical developmental trajectories. For instance, children with ADHD and other neurodevelopmental disorders often show atypical beta and theta activity during inhibitory tasks (Bocharov et al., 2021; Loo & Makeig, 2012; Michelini et al., 2022). The pronounced beta decrease in typically developing children observed here may provide a useful reference point for interpreting altered beta dynamics in clinical populations such as ADHD. These results underscore that a more detailed understanding of typical oscillatory maturation may facilitate the early identification and intervention for inhibitory control deficits.

### 4.5. Methodological considerations

This study leveraged the high temporal and spatial resolution of MEG and beamformer-based source modeling to identify temporal and spectral patterns of neural oscillatory dynamics involved in passive listening and response inhibition in children and adults. By focusing on modulations in alpha and beta power, we captured aspects of neural processing that are not phase-locked to stimulus onset and therefore remain unaddressed in traditional analyses of evoked responses. However, the relatively small adult sample limits the generalizability of the child–adult comparisons, which were therefore interpreted with caution. Moreover, because the child and adult cohorts are separated by an unsampled age range (15–19 years) and the child cohort is concentrated in middle childhood (87% aged 8–11 years; only three children younger than 8 and six aged 12 or older), the continuous-age models reported here cannot establish a continuous developmental trajectory.

A further methodological consideration concerns motor confounds. Electromyography (EMG) was not recorded during the MEG sessions, so covert or partial motor preparation that did not result in an overt button press could not be tracked at the trial level. Because the deviant tones analyzed here did not require an overt motor response in either task (withheld in Go/Nogo, absent in passive listening), epochs were matched for overt motor output across tasks. Go/Nogo epochs containing incorrect responses (commission errors) were excluded. This does not, however, rule out differential covert motor preparation or inhibition-related motor activity between the two tasks. Future studies should incorporate EMG or other movement-monitoring measures to allow trial-level control for such confounds.

In addition, the present analysis focused exclusively on alpha and low-beta frequency bands due to their well-established roles in response inhibition. Future studies should extend this approach to include the theta (4–8 Hz) and gamma (30–100 Hz) bands, which are increasingly recognized as critical for executive function and sensory integration (e.g., Uhlhaas et al., 2010; Tan et al., 2024).

### 4.6. Conclusion

This study demonstrates a clear developmental shift in the neural dynamics underlying inhibitory control. While children and adults engaged largely similar cortical regions during passive listening, the introduction of inhibitory demands revealed spectral, temporal, and spatial differences in oscillatory responses. Children relied on a prolonged beta power decrease in somatomotor and temporo-parietal regions overlapping in time with the anticipated motor response as estimated from the go-trial reaction times. Adults, in turn, showed temporally precise increases in alpha–beta power within a left-lateralized fronto-temporo-occipital network that emerged only after the expected motor response.

These findings suggest that, across development, inhibitory control is supported by increasingly efficient and coordinated neural strategies. In childhood, inhibition appears to depend on broad, sensorimotor-driven decrease, potentially reflecting slower and more elaborated activation of the motor system. In adulthood, the shift toward synchronized alpha– beta dynamics in executive networks enables more targeted and automated inhibition, and the observed oscillatory modulation in the late time window is likely related to the stabilization, maintenance, or evaluative processes underlying inhibitory control.

Taken together, our results refine current models of cognitive control by showing that effective inhibition is not only determined by regional recruitment but by the maturation of temporally and spectrally coordinated oscillatory mechanisms. This developmental trajectory may also provide a valuable framework for identifying atypical patterns of inhibitory control in clinical populations.

## Credit author statement Author contributions

*Finnilä, E.*: Conceptualization, Formal analysis, Visualization, Writing – original draft, Writing – review & editing.

*Van Bijnen, S.:* Conceptualization, Data curation, Investigation, Methodology, Project administration, Writing – review & editing.

*Kujala, J.*: Methodology, Visualization, Supervision, Writing – review & editing. *Parviainen, T.*: Conceptualization, Funding acquisition, Project administration, Resources, Supervision, Writing – review & editing.

## Data availability statement

The data that support the findings of this study are not publicly available due to ethical restrictions related to participant confidentiality. Anonymized data may be made available from the corresponding author upon reasonable request and with appropriate ethical approval.

## Declarations of interest

The authors declare no competing interests.

## Acknowledgements and Funding Statement

We thank all participants for their valuable contribution to this study. This work was supported by the Pirkanmaa Wellbeing Services County, the Viljo Syrenius Fund of the South Ostrobothnia Regional Fund of the Finnish Cultural Foundation (grant agreement no. 10241774), EU project ChildBrain (Horizon 2020 Marie Skłodowska-Curie Action (MSCA) Innovative Training Network (ITN) – European Training Network (ETN), grant agreement no. 641652) and Academy of Finland (grant agreement no. 370956).

---

**Supplementary Figure S1.**
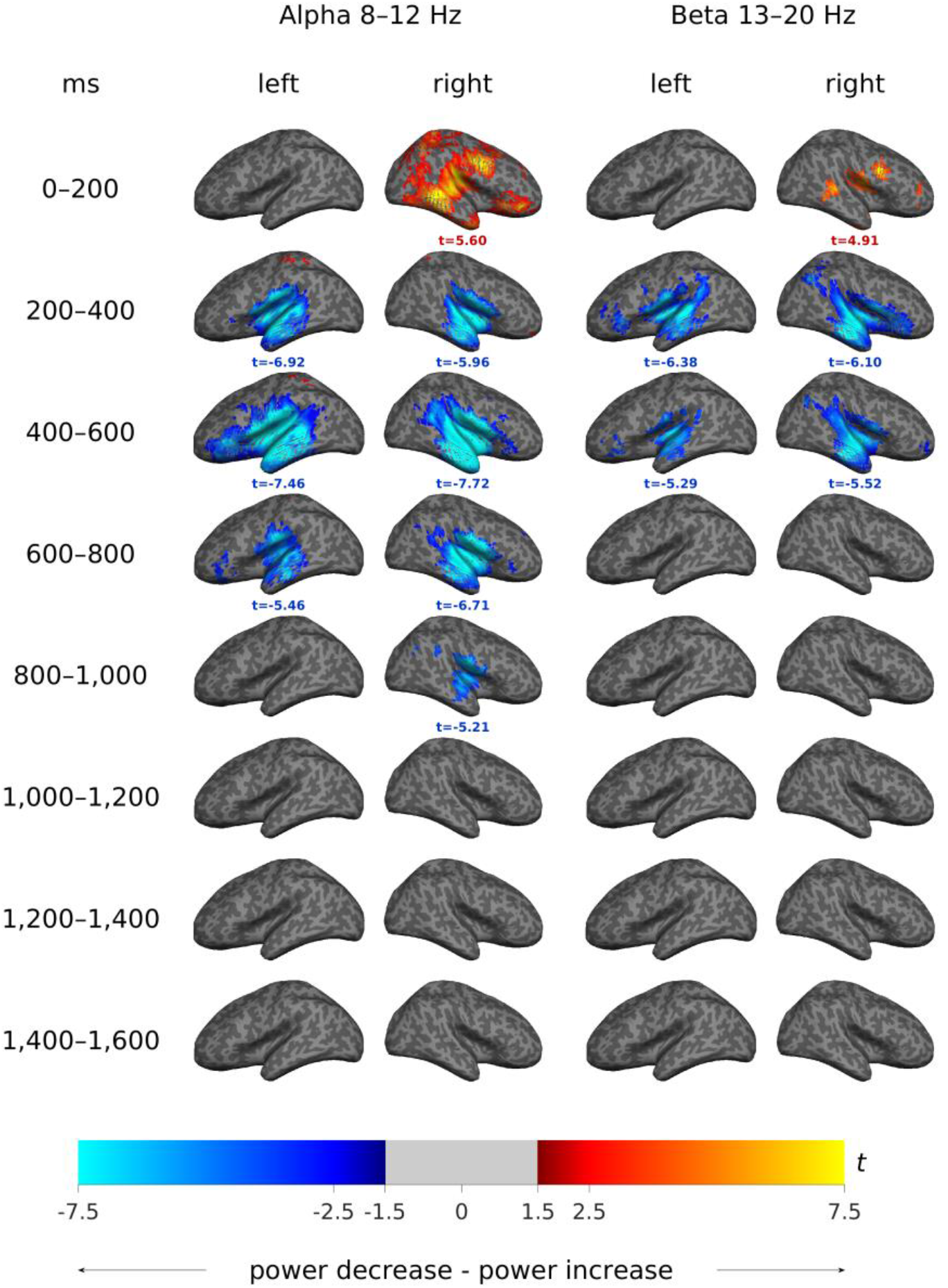
Whole-brain linear mixed-effects model, full sample: passive-listening baseline (intercept). Cortical surface maps show the model intercept — mean baseline-corrected alpha (8–12 Hz) and beta (13–20 Hz) power during passive listening at the sample mean age — thresholded at FDR q < .05 (corrected within each time window), across the full sample (children and adults combined, N = 83). Data are shown separately for the left and right hemispheres across eight consecutive 200-ms time windows (0–1,600 ms). Warm colors indicate power increases; cool colors indicate power decreases.

**Supplementary Figure S2.**
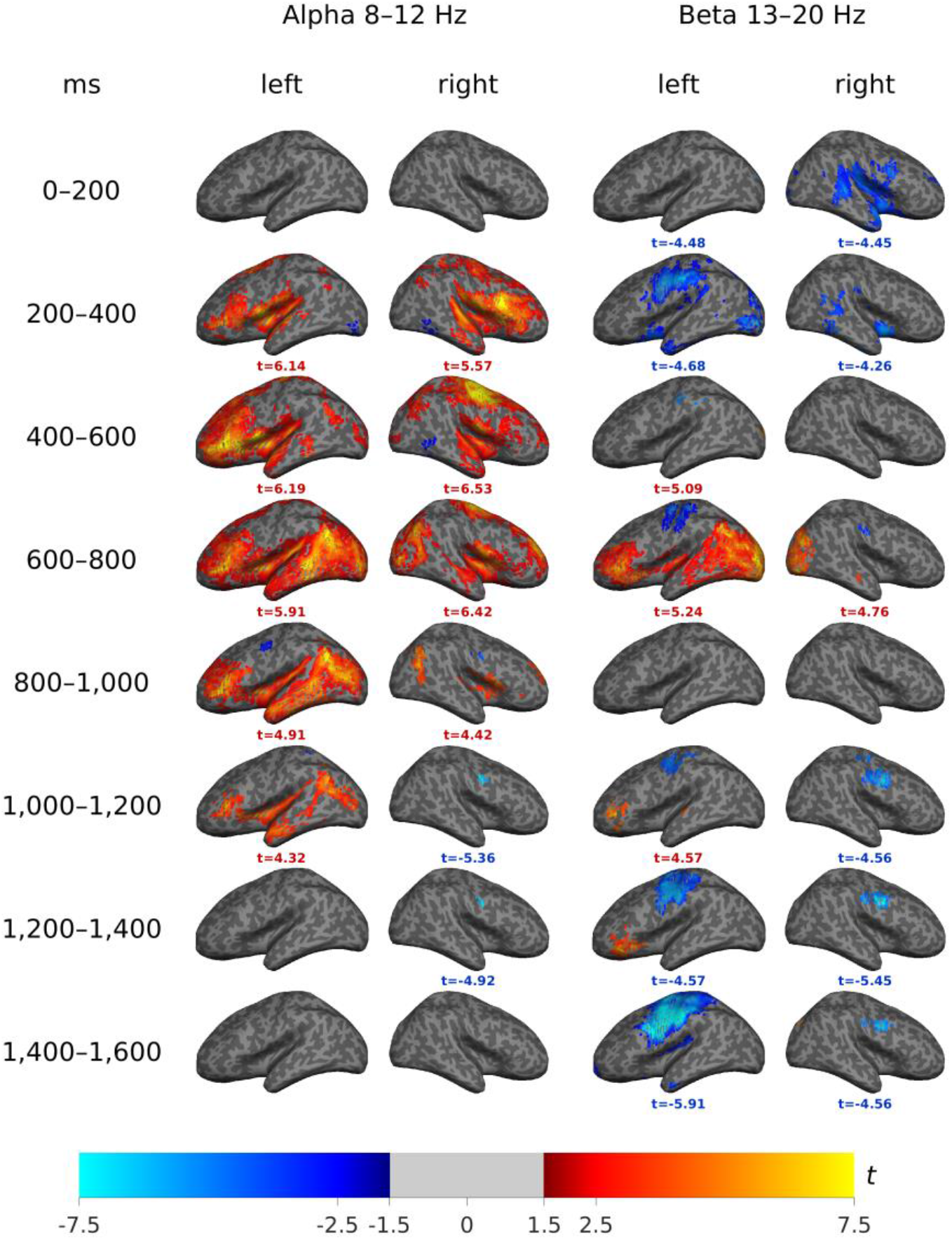
Whole-brain linear mixed-effects model, full sample: task effect (Go/Nogo minus passive). Cortical surface maps show the main effect of task, representing the Go/Nogo-minus-passive difference in oscillatory power at the sample mean age, across the full sample.

**Supplementary Figure S3.**
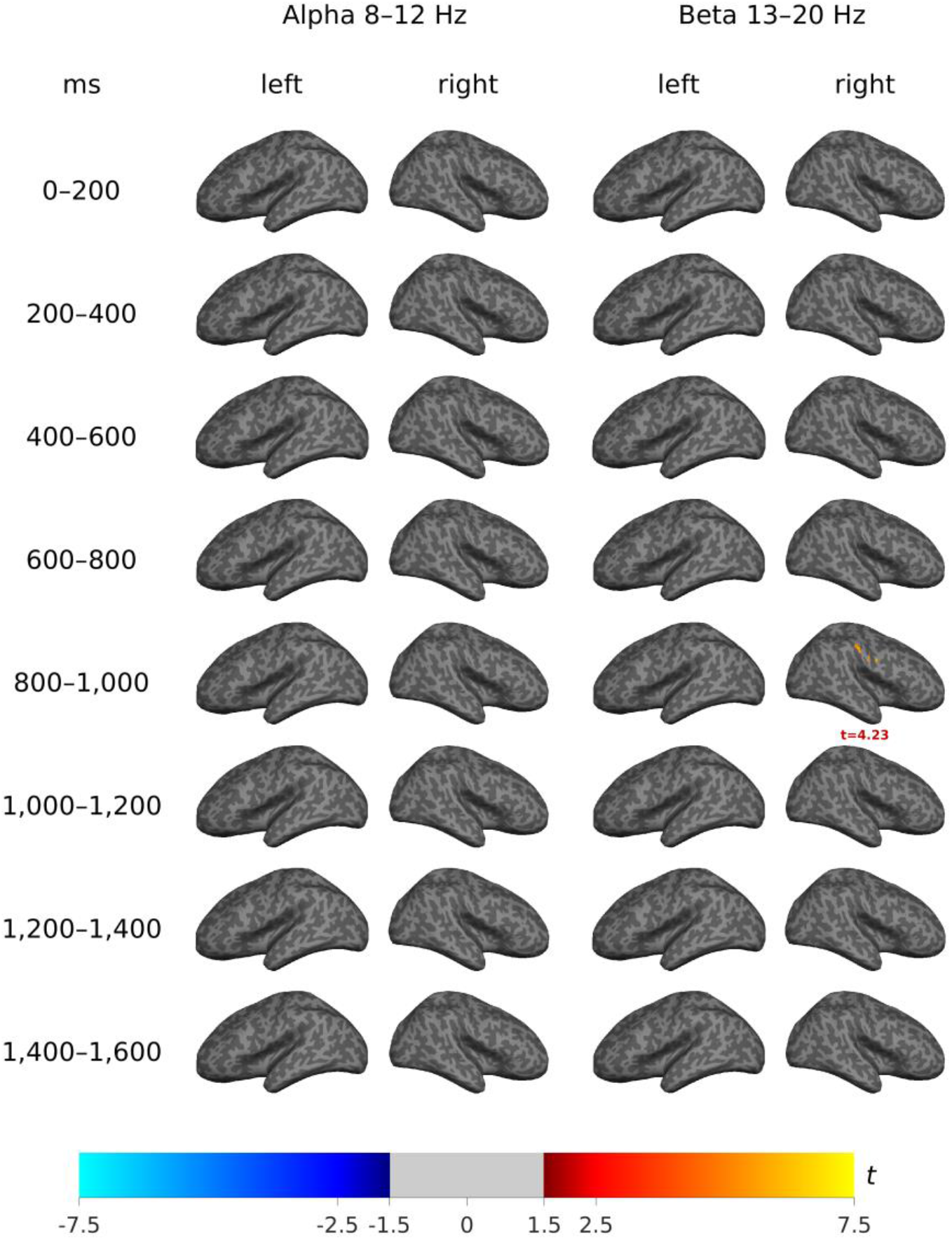
Whole-brain linear mixed-effects model, full sample: age effect during passive listening. Cortical surface maps show the main effect of age (linear age slope of baseline-corrected power during passive listening, given the task × age interaction in the model), across the full sample.

**Supplementary Figure S4.**
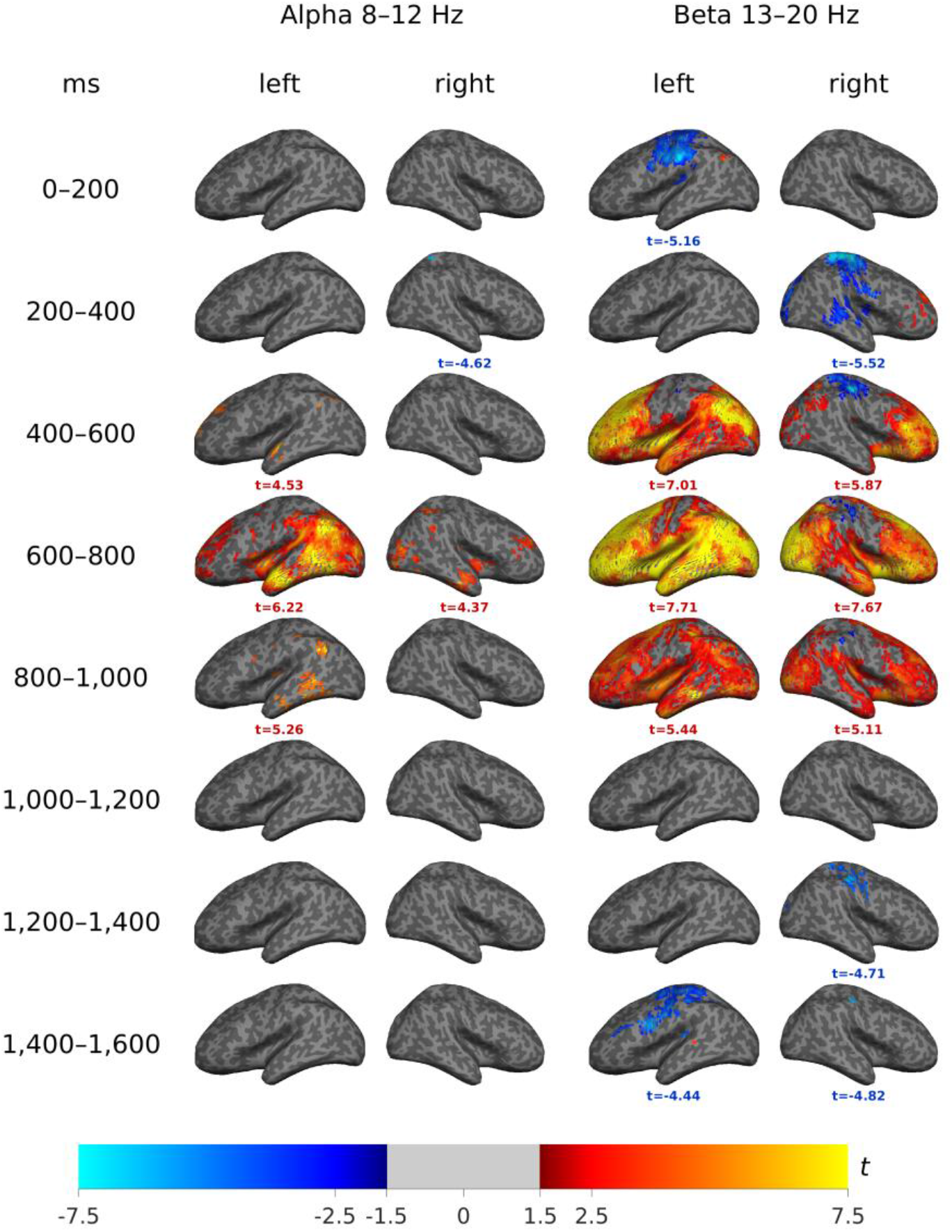
Whole-brain linear mixed-effects model, full sample: task × age interaction. Cortical surface maps show the task × age interaction, indexing how the Go/Nogo-minus-passive difference in oscillatory power changes with age across the full sample.

**Supplementary Figure S5.**
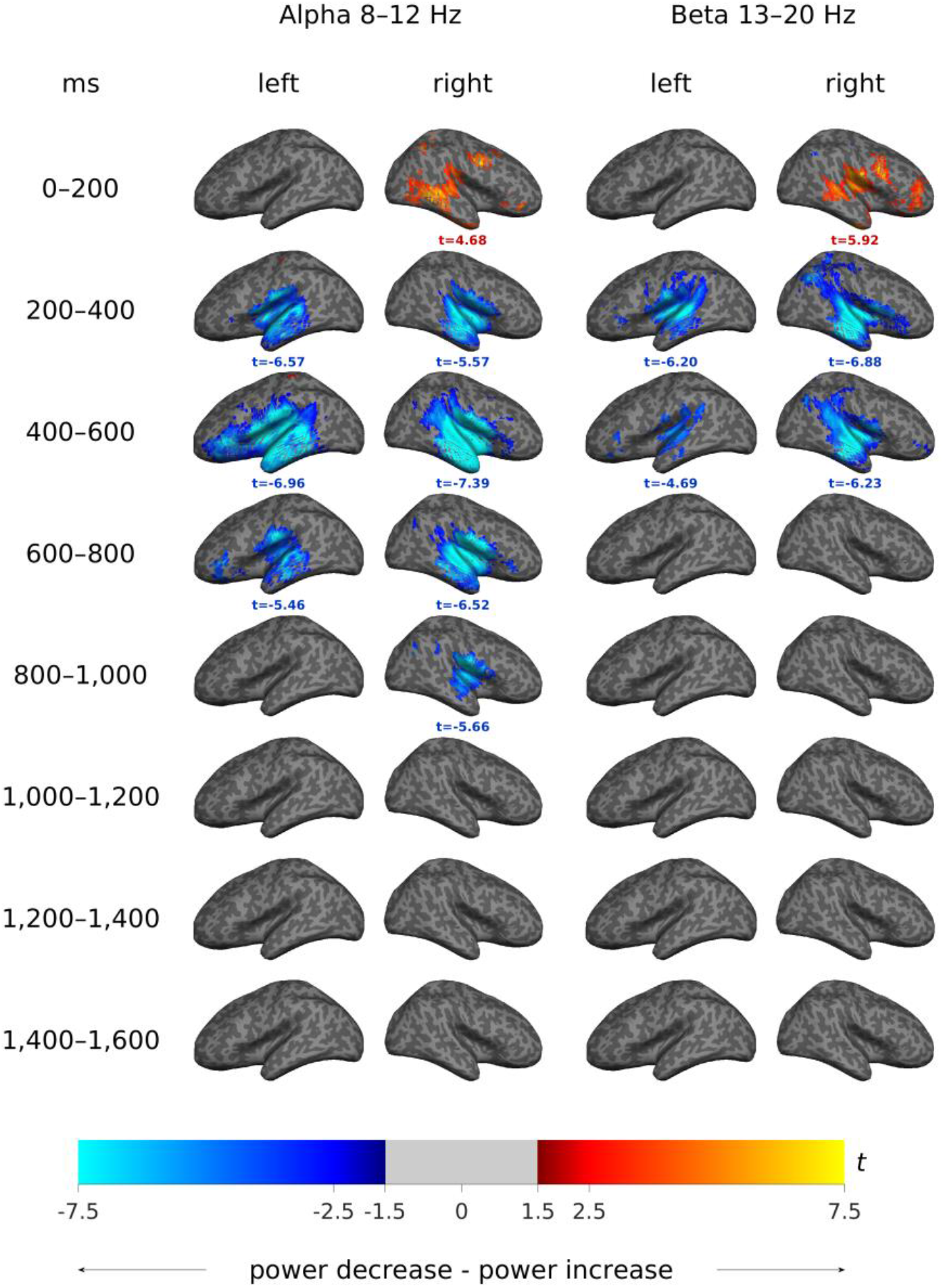
Whole-brain linear mixed-effects model, children only: passive-listening baseline (intercept). Cortical surface maps show the model intercept — mean baseline-corrected alpha (8–12 Hz) and beta (13–20 Hz) power during passive listening at the sample mean age — thresholded at FDR q < .05 (corrected within each time window), for the children’s cohort only (N = 67, age range 6–14 years). Data are shown separately for the left and right hemispheres across eight consecutive 200-ms time windows (0–1,600 ms). Warm colors indicate power increases; cool colors indicate power decreases.

**Supplementary Figure S6.**
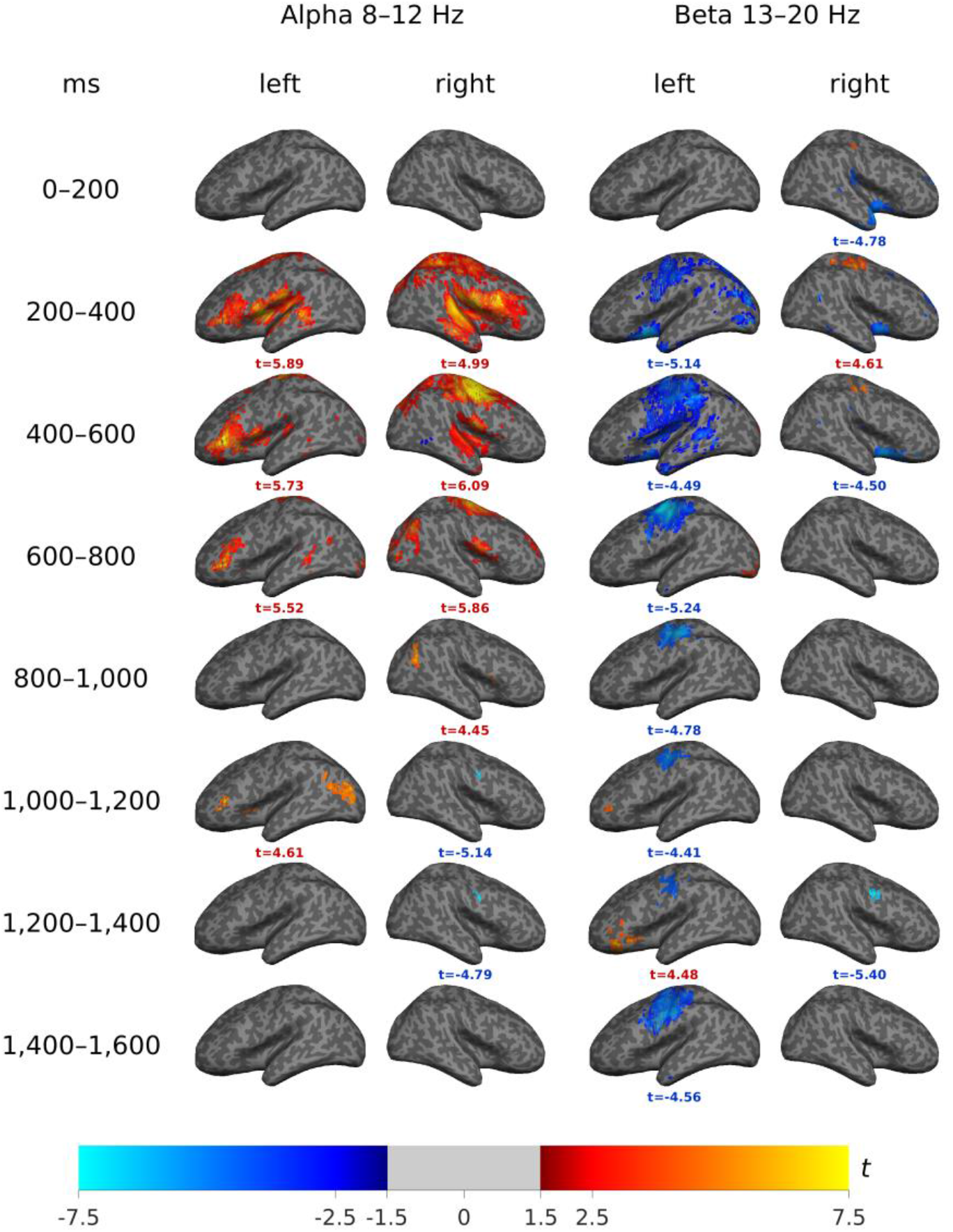
Whole-brain linear mixed-effects model, children only: task effect (Go/Nogo minus passive). Cortical surface maps show the main effect of task, representing the Go/Nogo-minus-passive difference in oscillatory power at the sample mean age, within the children’s cohort.

**Supplementary Figure S7.**
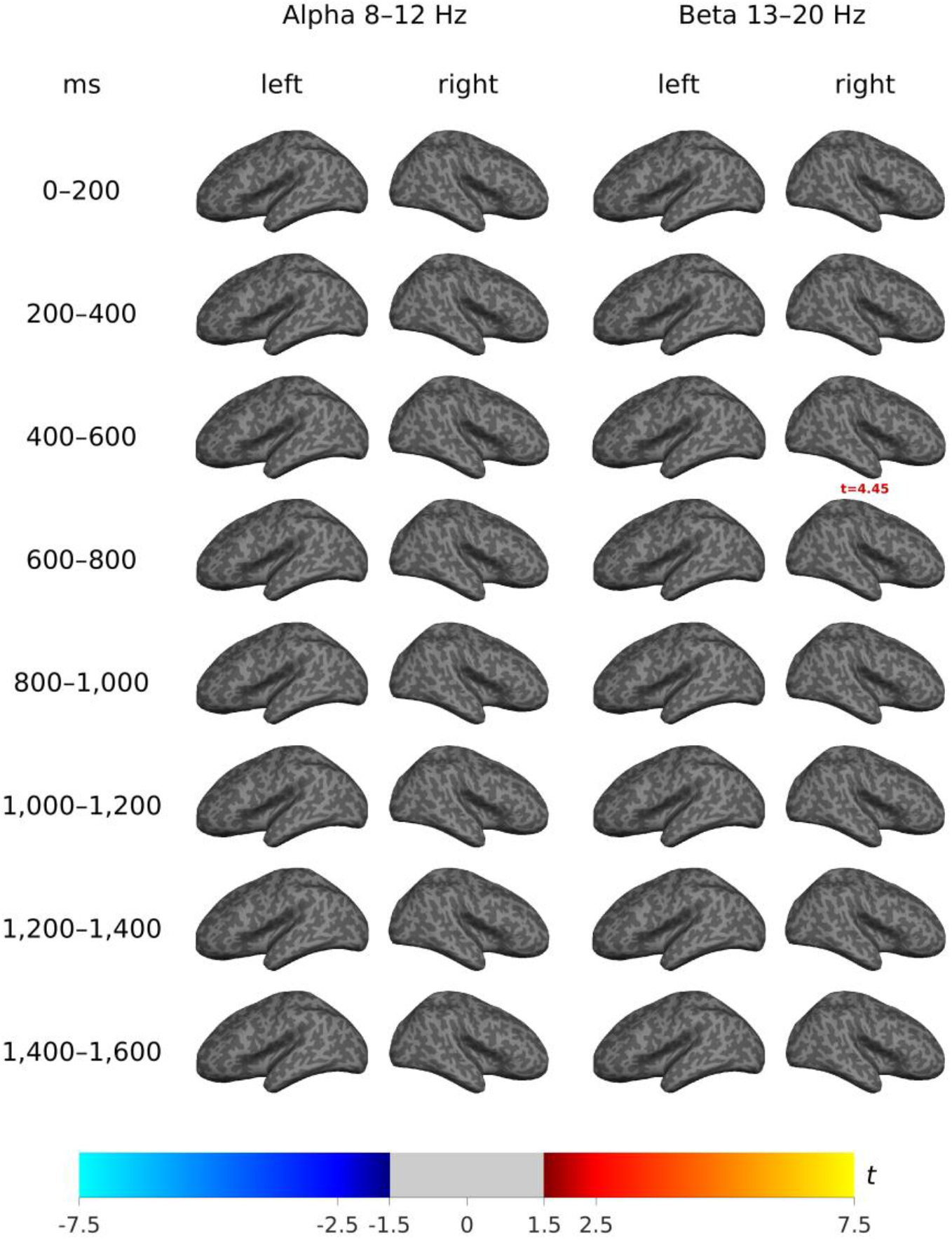
Whole-brain linear mixed-effects model, children only: age effect during passive listening. Cortical surface maps show the main effect of age (linear age slope of baseline-corrected power during passive listening, given the task × age interaction in the model) within the children’s cohort.

**Supplementary Figure S8.**
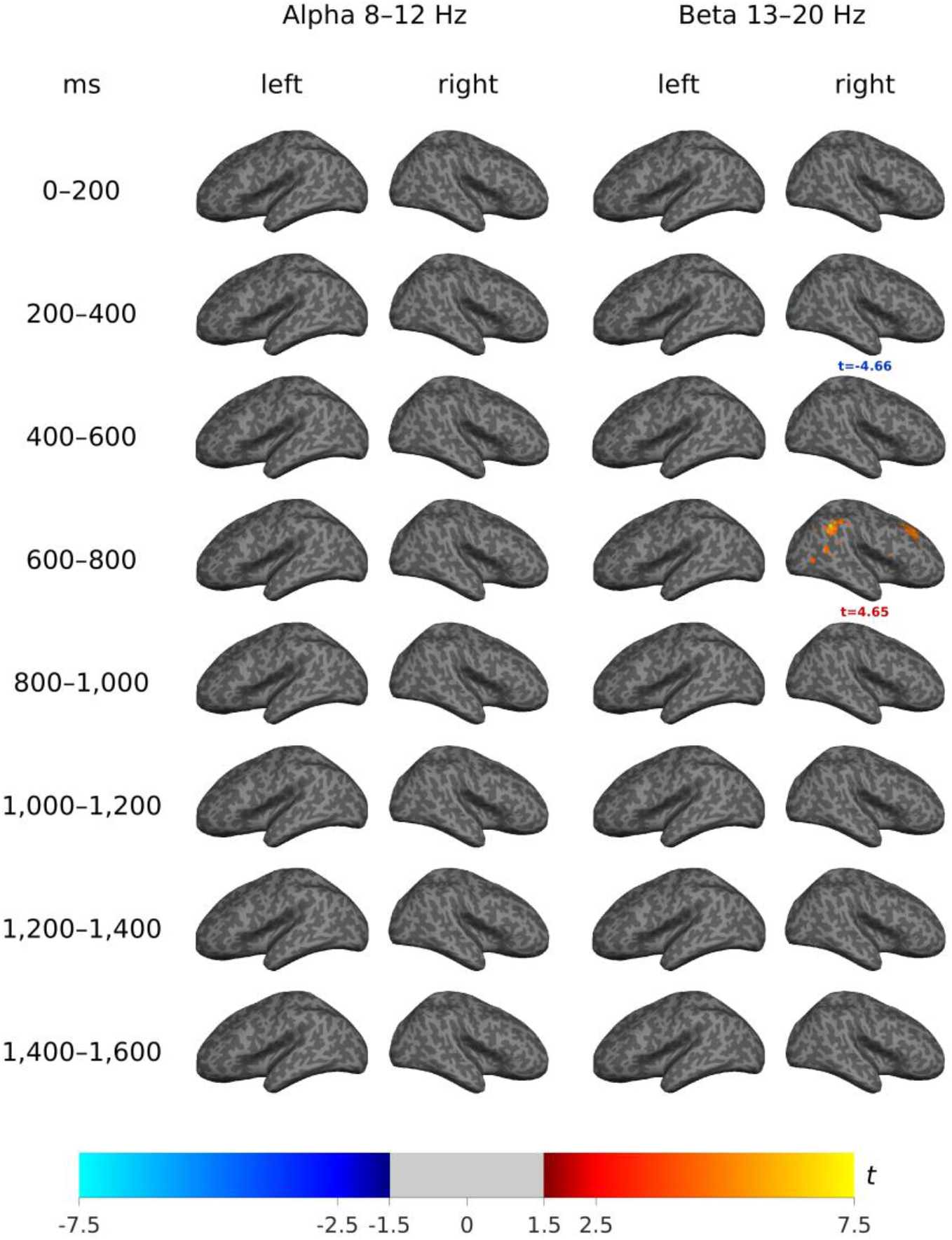
Whole-brain linear mixed-effects model, children only: task × age interaction. Cortical surface maps show the task × age interaction within the children’s cohort, indexing how the Go/Nogo-minus-passive difference in oscillatory power changes with age across childhood (6–14 years), independent of the child–adult group contrast.

**Supplementary Table S1.** FDR-significant clusters across the six whole-brain contrasts. Each row summarizes one FDR-significant cluster (hemisphere × frequency band × 200-ms time window, 0–1,600 ms post-stimulus) within the whole-brain contrast shown in the corresponding manuscript figure. Alpha = 8–12 Hz; Beta = 13–20 Hz. Peak t is the most extreme t-value among the significant vertices in that cluster, and its sign gives the direction of the effect; N vertices is the number of FDR-significant vertices; % positive is the proportion of those vertices with a positive t-value (together with the sign of Peak t, this identifies clusters with a substantial minority of oppositely signed vertices); p (FDR) is the smallest FDR-corrected p-value within the cluster. Some clusters consist of isolated suprathreshold vertices (N vertices = 1); these are reported for completeness but should be interpreted with caution. The Figure 4 section of this table lists only Go/Nogo clusters, as no significant age-group differences were found during passive listening (Section 3.3.1); this sixth contrast therefore contributes no rows.

| Age Group | Hemisphere | Band | Window (ms) | Peak *t* | *N* vertices | % positive | *p* (FDR) |
|---|---|---|---|---|---|---|---|
| **Figure 1 — Passive Listening Task (vs. Baseline)** |  |  |  |  |  |  |  |
| Adults | Left | Alpha | 400–600 | –6.18 | 154 | 4% | < .001 |
| Adults | Right | Alpha | 0–200 | 6.02 | 664 | 100% | < .001 |
| Children | Left | Alpha | 0–200 | 4.94 | 2 | 100% | < .001 |
| Children | Left | Alpha | 200–400 | –6.02 | 384 | 0% | < .001 |
| Children | Left | Alpha | 400–600 | –7.96 | 1,410 | 1% | < .001 |
| Children | Left | Alpha | 600–800 | –5.51 | 363 | 0% | < .001 |
| Children | Left | Beta | 200–400 | –6.19 | 778 | 0% | < .001 |
| Children | Left | Beta | 400–600 | –4.89 | 371 | 0% | < .001 |
| Children | Right | Alpha | 0–200 | 4.57 | 267 | 100% | < .001 |
| Children | Right | Alpha | 200–400 | –4.93 | 357 | 1% | < .001 |
| Children | Right | Alpha | 400–600 | –6.42 | 810 | 0% | < .001 |
| Children | Right | Alpha | 600–800 | –5.56 | 480 | 0% | < .001 |
| Children | Right | Alpha | 800–1,000 | –4.92 | 80 | 0% | < .001 |
| Children | Right | Beta | 0–200 | 5.76 | 280 | 100% | < .001 |
| Children | Right | Beta | 200–400 | –6.29 | 592 | 0% | < .001 |
| Children | Right | Beta | 400–600 | –5.72 | 505 | 0% | < .001 |
| Children | Right | Beta | 1,400–1,600 | –4.83 | 50 | 0% | < .001 |
| **Figure 2 — Go/Nogo Task (vs. Baseline)** |  |  |  |  |  |  |  |
| Adults | Left | Alpha | 400–600 | 5.10 | 126 | 100% | < .001 |
| Adults | Left | Alpha | 600–800 | 5.73 | 1,408 | 100% | < .001 |
| Adults | Left | Alpha | 800–1,000 | 5.34 | 969 | 100% | < .001 |
| Adults | Left | Beta | 600–800 | 5.69 | 1,588 | 100% | < .001 |
| Adults | Left | Beta | 800–1,000 | 6.53 | 1 | 100% | < .001 |
| Children | Left | Alpha | 0–200 | 6.69 | 1,709 | 100% | < .001 |
| Children | Left | Alpha | 200–400 | 6.12 | 1,006 | 84% | < .001 |
| Children | Left | Alpha | 400–600 | –6.07 | 1,219 | 40% | < .001 |
| Children | Left | Alpha | 600–800 | 5.23 | 695 | 68% | < .001 |
| Children | Left | Alpha | 1,000–1,200 | 5.18 | 162 | 100% | < .001 |
| Children | Left | Alpha | 1,200–1,400 | 4.94 | 1 | 100% | < .001 |
| Children | Left | Beta | 200–400 | –6.15 | 2,868 | 0% | < .001 |
| Children | Left | Beta | 400–600 | –7.20 | 2,522 | 1% | < .001 |
| Children | Left | Beta | 600–800 | –4.91 | 594 | 21% | < .001 |
| Children | Left | Beta | 800–1,000 | –4.62 | 245 | 0% | < .001 |
| Children | Left | Beta | 1,000–1,200 | –5.00 | 256 | 0% | < .001 |
| Children | Left | Beta | 1,200–1,400 | –4.86 | 319 | 0% | < .001 |
| Children | Left | Beta | 1,400–1,600 | –5.62 | 928 | 0% | < .001 |
| Children | Right | Alpha | 0–200 | 6.10 | 1,747 | 100% | < .001 |
| Children | Right | Alpha | 200–400 | 6.22 | 1,150 | 97% | < .001 |
| Children | Right | Alpha | 400–600 | 5.99 | 1,346 | 60% | < .001 |
| Children | Right | Alpha | 600–800 | –5.83 | 914 | 69% | < .001 |
| Children | Right | Alpha | 800–1,000 | 4.43 | 156 | 78% | < .001 |
| Children | Right | Beta | 200–400 | –7.53 | 1,414 | 3% | < .001 |
| Children | Right | Beta | 400–600 | –7.07 | 1,285 | 4% | < .001 |
| Children | Right | Beta | 600–800 | –4.48 | 294 | 15% | < .001 |
| Children | Right | Beta | 800–1,000 | –4.78 | 151 | 0% | < .001 |
| Children | Right | Beta | 1,000–1,200 | –4.64 | 179 | 0% | < .001 |
| Children | Right | Beta | 1,200–1,400 | –5.77 | 167 | 0% | < .001 |
| Children | Right | Beta | 1,400–1,600 | –4.93 | 221 | 0% | < .001 |
| **Figure 3 — Passive Listening vs. Go/Nogo (Task Contrast)** |  |  |  |  |  |  |  |
| Adults | Left | Alpha | 400–600 | 8.18 | 358 | 100% | < .001 |
| Adults | Left | Alpha | 600–800 | 8.43 | 1,557 | 100% | < .001 |
| Adults | Left | Alpha | 800–1,000 | 5.55 | 1,009 | 100% | < .001 |
| Adults | Left | Beta | 400–600 | 6.44 | 5 | 100% | < .001 |
| Adults | Left | Beta | 600–800 | 8.11 | 1,816 | 100% | < .001 |
| Adults | Right | Alpha | 400–600 | 7.40 | 103 | 100% | < .001 |
| Adults | Right | Alpha | 600–800 | 7.68 | 98 | 100% | < .001 |
| Children | Left | Alpha | 200–400 | 5.84 | 819 | 100% | < .001 |
| Children | Left | Alpha | 400–600 | 5.65 | 553 | 100% | < .001 |
| Children | Left | Alpha | 600–800 | 5.22 | 311 | 100% | < .001 |
| Children | Left | Beta | 200–400 | –4.91 | 380 | 0% | < .001 |
| Children | Left | Beta | 400–600 | –4.42 | 726 | 2% | < .001 |
| Children | Left | Beta | 600–800 | –4.93 | 358 | 20% | < .001 |
| Children | Left | Beta | 800–1,000 | –4.70 | 130 | 0% | < .001 |
| Children | Left | Beta | 1,400–1,600 | –4.37 | 271 | 0% | < .001 |
| Children | Right | Alpha | 200–400 | 4.65 | 961 | 100% | < .001 |
| Children | Right | Alpha | 400–600 | 5.68 | 752 | 100% | < .001 |
| Children | Right | Alpha | 600–800 | 4.94 | 403 | 100% | < .001 |
| Children | Right | Alpha | 1,000–1,200 | –4.93 | 1 | 0% | < .001 |
| Children | Right | Alpha | 1,200–1,400 | –4.58 | 5 | 0% | < .001 |
| Children | Right | Beta | 200–400 | –4.32 | 77 | 36% | < .001 |
| Children | Right | Beta | 1,200–1,400 | –5.04 | 12 | 0% | < .001 |
| **Figure 4 — Age-Group Differences, Go/Nogo Task** |  |  |  |  |  |  |  |
| — | Left | Alpha | 400–600 | –5.61 | 412 | 0% | < .001 |
| — | Left | Alpha | 600–800 | –5.64 | 1,254 | 0% | < .001 |
| — | Left | Beta | 400–600 | –6.87 | 1,417 | 0% | < .001 |
| — | Left | Beta | 600–800 | –6.26 | 2,245 | 0% | < .001 |
| — | Left | Beta | 800–1,000 | –5.41 | 1 | 0% | < .001 |
| **Figure 5 — Age-Group Differences, Task Contrast** |  |  |  |  |  |  |  |
| — | Left | Alpha | 400–600 | –5.34 | 71 | 0% | < .001 |
| — | Left | Alpha | 600–800 | –5.62 | 301 | 0% | < .001 |
| — | Left | Beta | 400–600 | –5.46 | 1,115 | 0% | < .001 |
| — | Left | Beta | 600–800 | –6.03 | 2,265 | 0% | < .001 |

**Supplementary Table S2.** Cortical parcels ranked by overall oscillatory power modulation relative to baseline (30 strongest parcels). For each Desikan–Killiany parcel and hemisphere, baseline-corrected power was averaged across the parcel’s source points, and the effect size of the modulation relative to baseline (Cohen’s d = mean / SD across participants) was computed separately for each group, task, frequency band, and time window. The score (Mean |d|) is the maximum over time windows (0–1,600 ms) of the absolute d averaged across both tasks (passive listening and Go/Nogo), both frequency bands (alpha, beta), and both age groups (children and adults); effect sizes were used so that the score is not biased by the unequal group sizes. Parcels are ranked in descending order of this score. Three ROIs representing the two most prominently modulated, spatially distinct clusters, the left and right lateral temporal cortex and the left inferior frontal gyrus, were selected for the time-course analyses shown in Figure 6.

| Rank | Hemisphere | Region | Mean \|d\| | ROI |
|---|---|---|---|---|
| **All conditions combined (vs. Baseline)** |  |  |  |  |
| 1 | Right | Superior temporal | 0.635 | LTC |
| 2 | Left | Superior temporal | 0.622 | LTC |
| 3 | Left | Transverse temporal | 0.616 | LTC |
| 4 | Right | Insula | 0.597 |  |
| 5 | Right | Transverse temporal | 0.589 | LTC |
| 6 | Left | Insula | 0.583 |  |
| 7 | Right | Middle temporal | 0.559 | LTC |
| 8 | Left | Middle temporal | 0.495 | LTC |
| 9 | Left | Supramarginal | 0.493 |  |
| 10 | Right | Parahippocampal | 0.487 |  |
| 11 | Left | Parahippocampal | 0.482 |  |
| 12 | Left | Pars triangularis | 0.461 | IFG |
| 13 | Right | Rostral middle frontal | 0.440 |  |
| 14 | Left | Inferior temporal | 0.440 |  |
| 15 | Left | Pars orbitalis | 0.436 | IFG |
| 16 | Left | Postcentral | 0.431 |  |
| 17 | Left | Pars opercularis | 0.428 | IFG |
| 18 | Right | Postcentral | 0.428 |  |
| 19 | Left | Fusiform | 0.423 |  |
| 20 | Right | Pars triangularis | 0.422 |  |
| 21 | Left | Rostral middle frontal | 0.420 |  |
| 22 | Right | Supramarginal | 0.416 |  |
| 23 | Left | Superior frontal | 0.413 |  |
| 24 | Right | Banks of STS | 0.412 | LTC |
| 25 | Left | Lingual | 0.406 |  |
| 26 | Right | Frontal pole | 0.404 |  |
| 27 | Left | Banks of STS | 0.403 | LTC |
| 28 | Left | Inferior parietal | 0.402 |  |
| 29 | Right | Superior parietal | 0.396 |  |
| 30 | Left | Precentral | 0.393 |  |

## Notes

### Competing Interest Statement

The authors have declared no competing interest.

